# Emerging role of oncogenic β-catenin in exosome biogenesis as a driver of immune escape in hepatocellular carcinoma

**DOI:** 10.1101/2023.10.24.563805

**Authors:** Camille Dantzer, Justine Vaché, Aude Brunel, Isabelle Mahouche, Anne-Aurélie Raymond, Jean-William Dupuy, Melina Petrel, Paulette Bioulac-Sage, David Perrais, Nathalie Dugot-Senant, Mireille Verdier, Barbara Bessette, Clotilde Billottet, Violaine Moreau

## Abstract

Immune checkpoint inhibitors have produced encouraging results in cancer patients. However, the majority of β-catenin-mutated tumors have been described as lacking immune infiltrates and resistant to immunotherapy. The mechanisms by which oncogenic β-catenin affects immune surveillance remain unclear. Herein, we highlighted the involvement of β-catenin in the regulation of the exosomal pathway and, by extension, in immune/cancer cell communication in hepatocellular carcinoma (HCC). We showed that mutated β-catenin represses expression of *SDC4* and *RAB27A*, two main actors in exosome biogenesis, in both liver cancer cell lines and HCC patient samples. Using nanoparticle tracking analysis and live-cell imaging, we further demonstrated that activated β-catenin represses exosome release. Then, we demonstrated in 3D spheroid models that activation of β-catenin promotes a decrease in immune cell infiltration through a defect in exosome secretion. Taken together, our results provide the first evidence that oncogenic β-catenin plays a key role in exosome biogenesis. Our study gives new insight into the impact of β-catenin mutations on tumor microenvironment remodeling, which could lead to the development of new strategies to enhance immunotherapeutic response.

## INTRODUCTION

Hepatocellular carcinoma (HCC) is the most frequent form of primary liver adult cancer (80% of cases). HCC represents the 6^th^ most common diagnosed cancer worldwide and the 3^rd^ leading cause of cancer-related death (1). This pathology has a poor prognosis and the diagnosis is often made at an advanced stage for which treatment options are limited, especially if surgery is no longer possible. In 2020, the combination of Atezolizumab (anti-PD-L1) and Bevacizumab (anti-VEGF) became the first-line FDA-approved therapy for advanced HCC (2). Despite this therapeutic advance, several studies emphasized that β-catenin-mutated HCC are devoid of immune infiltrates and are resistant to immunotherapy (3–6). *CTNNB1* gene mutations are found in 30-40% of HCC and trigger uncontrolled transcriptional activity (7,8). These mutations prevent β-catenin degradation, fostering its role as a transcriptional co-factor and thus the expression of genes involved in cell proliferation and invasion. This altered pattern of gene expression could provide a key to understanding to the observed immune evasion. β-catenin, encoded by the *CTNNB1* gene, is a main oncogene, mutated in various cancers, such as melanoma and liver, endometrial, and colorectal cancers (9). These β-catenin-mutated tumors share specific features, prominent among them a microenvironment devoid of immune infiltrates. Thus, despite of a therapeutic revolution for cancer treatment, with the emergence of immunotherapy and more particularly immune checkpoint inhibitors (anti-PD1, anti-PD-L1), most of the β-catenin-mutated tumors remain resistant to immunotherapy (10–15). In these tumors, the oncogenic β-catenin is able to establish a microenvironment that favors tumor progression notably by promoting immune escape. Few studies have reported that the oncogenic β-catenin is implicated in the impairment of intercellular communication between cancer cells and immune cells, partly through soluble molecules such as cytokines.. In melanoma and HCC, a decrease in CCL4 and CCL5, leads to a defective recruitment of dendritic cells and consequently impaired T-cell activity and immune escape (5,11). As immunotherapies are now part of the growing arsenal for treating cancers, it is important to better understand the mechanisms underlying this immune escape phenotype. In this study we focused on β-catenin-mutated HCC and intercellular communication through extracellular vesicles.

Tumor-derived extracellular vesicles (EVs) are increasingly described as important actors in tumor microenvironment communication (16); but no studies explored their role in the immune escape and immunotherapy resistance of β-catenin-mutated tumors. EVs are nanometric phospholipid bilayer transport vesicles, which contain various cargoes such as RNAs or proteins and that impact the properties and functions of recipient cells. EVs can be classified into two main categories: microvesicles and exosomes. Microvesicles (50 to 1000 nm) correspond to EVs directly secreted from the plasma membrane (PM). By contrats, exosomes (50 to 150 nm) are released from an endosomal compartment, the multivesicular body (MVB), by its fusion with the cell surface (16). Among the actors implicated in exosome biogenesis and secretion, syndecan-4, involved in the endosomal membrane budding (17) and Rab27a, which promotes the docking of MVB to the PM, play key roles in exosome release (18). As important mediators of cell-to-cell communication, EVs can regulate tumor growth, angiogenesis, invasion and infiltration of immune cells into the tumor (19). In the context of HCC, it was reported that EVs derived from tumor cells can modulate epithelial-to-mesenchymal transition, intrahepatic metastasis and tumor immunity (20).

Based on these considerations, the current study investigated the role of exosomes in mutated β-catenin-mediated immune evasion in HCC. We demonstrated that mutated β-catenin represses syndecan-4 and Rab27a expression, thus altering exosomal secretion from HCC cells, in turn leading to a defective recruitment of immune cells in tumors.

## RESULTS

### Silencing of mutated β-catenin increases exosome biogenesis-associated gene expression in HepG2 cells

*CTNNB1* mutations in HCC are often monoallelic, leaving a wild-type allele in tumor cells. To study the oncogenic β-catenin specifically, we made use of the dual β- catenin knockdown (KD) HepG2 model we published recently (21). The transcriptional analysis of HepG2 KD cells revealed that the expression of 1973 genes (log2(Fold-change) >0.5 and <-0.5) is modulated upon mutated β-catenin silencing (21) (Figure 1a). Gene ontology analysis (FunRich) performed on upregulated genes showed that mutated β-catenin silencing enhanced significantly the expression of genes linked to specific cellular components, including exosomes (Figure 1b). Moreover, silencing of the mutated β-catenin led to the overexpression of 17 genes associated with exosome biogenesis, such as *RAB27A* and *SDC4* (Figure 1c). Similarly, proteomic analysis of HepG2 KD cells revealed exosomes as the main cellular component displaying up-regulated proteins upon mutated β- catenin silencing (Figures 1d-1e). Consistent with the transcriptomic analysis, we found Rab27a and Syndecan-4 among up-regulated proteins, results (Figure 1f), even if the variability was found for Syndecan-4 protein levels. These results suggest the involvement of mutated β-catenin in the regulation of the exosomal pathway.

**Figure 1.**
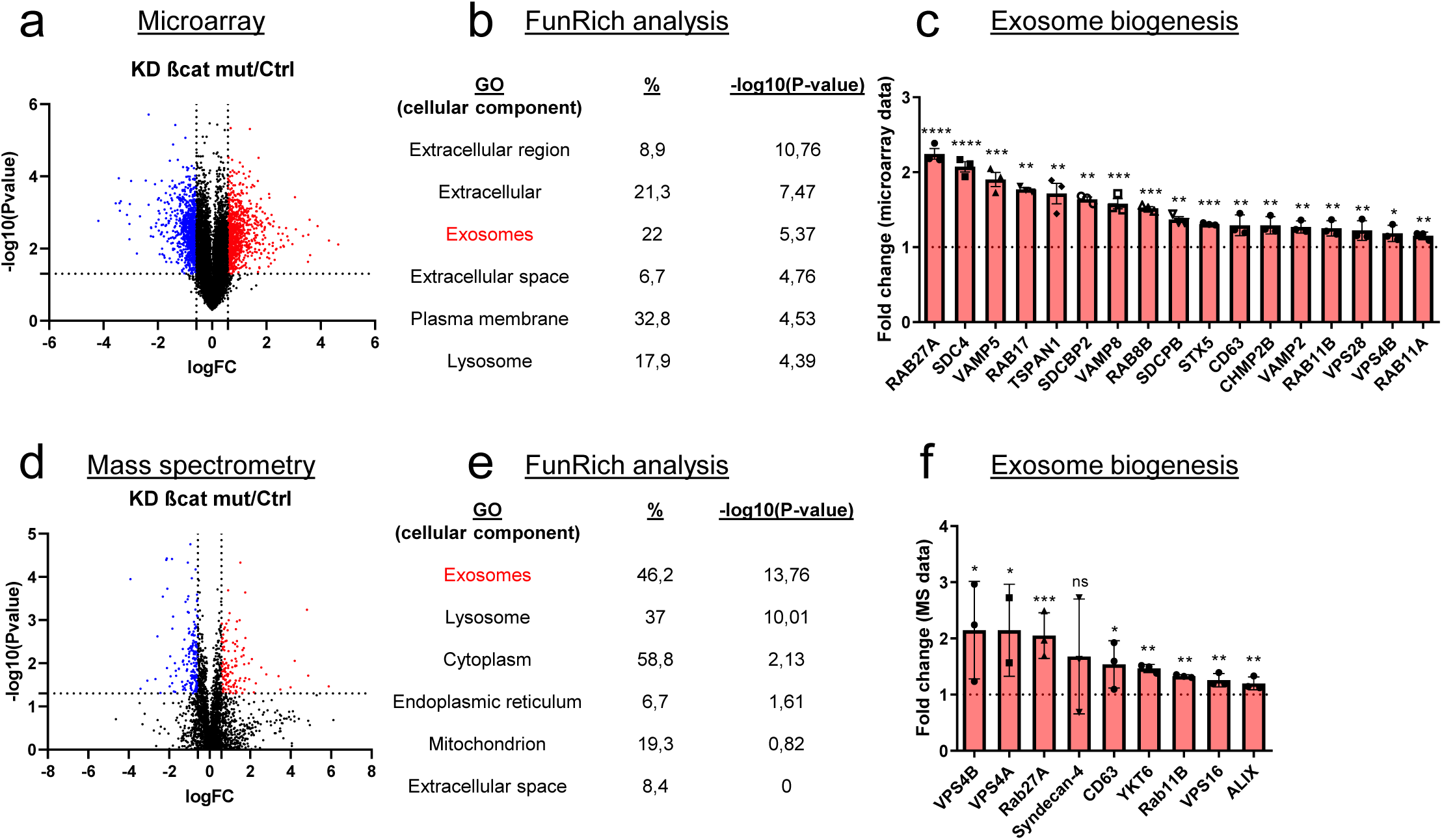
Mutated β-catenin regulates exosome biogenesis gene expression in HepG2 cells. (**a-f**) HepG2 cells were transfected with siRNAs targeting mutated β- catenin or control siRNAs. **(a)** Volcano plot of deregulated genes identified by microarray based-transcriptomic analysis. Red and blue dots indicated respectively significantly up- and down-regulated genes. **(b)** Upregulation of cellular component genes using FunRich software. **(c)** Upregulated genes associated with exosome biogenesis pathway. The graph indicates the fold change (FC) when comparing the mutated β-catenin silencing condition with the control condition. **(d)** Volcano plot of deregulated proteins identified by mass spectrometry. Red and blue dots indicated respectively significantly up- and down-regulated proteins. **(e)** Upregulation of cellular component proteins identified using FunRich software. **(f)** Upregulated proteins associated with exosome biogenesis pathway. The graph indicates the fold change when comparing the mutated β-catenin silencing condition with the control condition. Results are expressed as Mean ± SEM, two-tailed Student’s t-test analysis. *P<0.05;**P<0.01; ***P<0.001; ****P<0.0001.

### Stable depletion of mutated β-catenin increases exosome secretion in the HepG2 cell model

In the model presented above, siRNAs were used to transiently silence mutated β- catenin expression in HepG2 cells (21). To improve this method, the same sequences were used to develop an inducible shRNA strategy to stably inhibit the expression of the mutated β-catenin. Through the use of doxycycline-inducible promoters we were able to regulate the expression of shRNAs over time for more in-depth and specific investigations of β-catenin function in various cellular processes. In HepG2 cells, this strategy reduced protein expression by 62% for mutated β- catenin and by 42% for CyclinD1, a β-catenin-positive transcriptional target (Figure 2a). The model was also validated regarding mRNA expression of several β-catenin transcriptional targets. We noted a decrease in the expression of *CCND1* and *AXIN2* (positive targets) and an increase in the expression of *ARG1* expression (negative target) (Figure sup 1a). Also, as previously described with the siRNA strategy (21), after mutated β-catenin depletion in HepG2 cells, we observed an increase in the number and size of bile canaliculi (BC), feature of more differentiated cells (Figures sup 1b-1c). Nanoparticle-tracking analysis (NTA) revealed that depleting mutated β- catenin using either siRNA or shRNA increased the number of secreted particles (Figure 2b, Figure sup 1d). The particle mean size (100-150 nm), unaltered by either approach, was consistent with the size of typical small EVs, such as exosomes (Figure 2b, Figure sup 1d). To analyze the exosomal release in living HepG2 cells, we used a pH-sensitive reporter (CD63-pHluorin) that allowed the visualization of MVB–PM fusion by total internal reflection fluorescence (TIRF) microscopy (22). The tetraspanin CD63 is a well-known exosome surface marker (23,24). In cells expressing CD63-pHluorin, we could detect multiple discrete increases in the fluorescence signal at the PM over the time suggesting ongoing fusion of CD63-pHluorin–positive acidic vesicles with the PM (Figure sup 1e). We found that mutated β-catenin depletion in HepG2 cells significantly increased the number of fusion events associated with secreted exosomes (Figure 2c). EVs released by HepG2 cells were next isolated by differential ultracentrifugation and characterized (Figure 2d). First, western-blot analysis of isolated EVs confirmed the expression of the CD63 exosomal marker. It is noteworthy that we observed an increase in CD63 expression after mutated β-catenin depletion, suggesting an overall increase in the exosomal fraction (Figure 2e). This increased expression of CD63 was also confirmed with the proteomic analysis of HepG2 KD cells (Figure 1f). Moreover, the WT and the mutated forms of the β-catenin protein were also found in HepG2-derived EVs, and the expression of mutated β-catenin was decreased relative to the cells of origin (Figure 2e). Using transmission electron microscopy (TEM), we demonstrated that HepG2- derived EVs have the morphology (flat cup-shaped) and size (50-150nm) of typical (25) (Figures 2f and sup 1f). To better understand the regulation of exosome secretion by mutated β-catenin, we next used TEM to visualize MVBs, the origin of exosomes. We observed a higher number of MVBs per cell and an increase in their diameter upon removal of mutated β-catenin (Figure 2g). Taken together these results highlight that mutated β-catenin represses exosome secretion.

**Figure 2.**
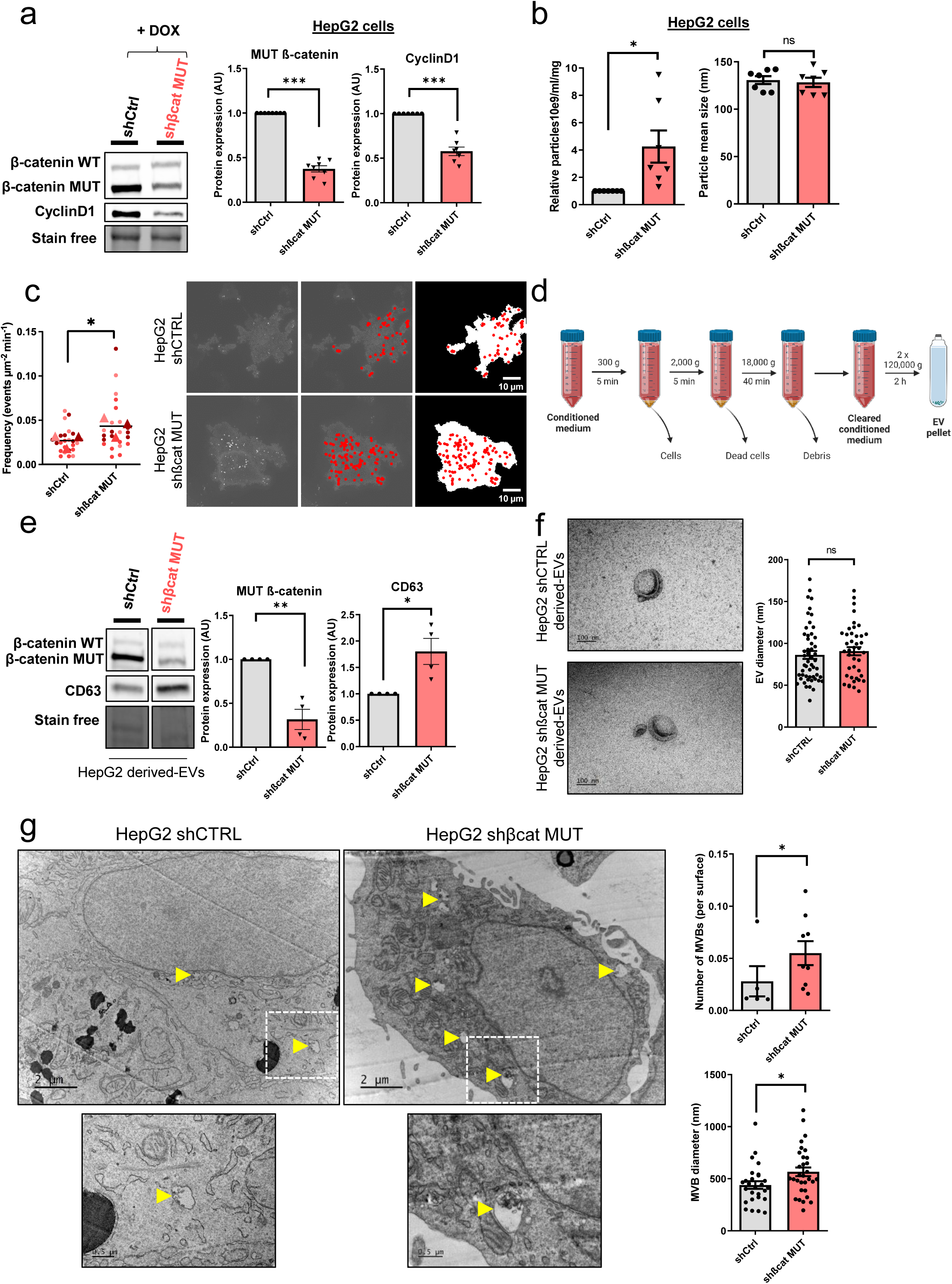
Mutated β-catenin controls exosome secretion in HepG2 cells. (**a-g**) HepG2-shβCat MUT and HepG2-shCtrl cells were treated with doxycycline (DOX) to silence or not mutated β-catenin. **(a)** Analysis of β-catenin and CyclinD1 expression by western-blot. Stain free was used as loading control. Graphs show the quantification of seven independent experiments. **(b)** Nanoparticle tracking analysis of supernatant. Graphs show the quantification of seven independent experiments. **(c)** Quantification of CD63-pHluorin MVB–PM fusion events visualized by live TIRF microscopy. Depicted data are representative of three independent experiments, each dot represents one cell. Images represent the cell mask (white) and red dots corresponding to fusion events. Scale bar: 10µm. **(d)** Extracellular vesicles (EVs) isolation protocol. Created with BioRender.com. **(e)** Analysis of β-catenin and CD63 expression in HepG2-derived EVs. Stain free was used as loading control. Graphs show the quantification of four independent experiments. **(f)** Transmission electron microscopy images of HepG2-derived EVs by close-up. Scale bar: 100nm. The graph shows the diameter quantification of EVs (n=93). **(g)** Electron microscopy images of HepG2 shCtrl and shβcat MUT cells showing multivesicular bodies (MVBs) (yellow arrowheads). Scale bar: 2 µm (zoom: 500 nm). The graphs show the quantification of the number of MVBs per cell and the MVB diameter. (**a-f**) Results are expressed as Mean ± SEM, two-tailed Student’s t-test analysis. *P<0.05; **P<0.01; ***P<0.001; ns, non-significant.

### Mutated β-catenin decreases Syndecan-4 and Rab27a expression in liver cancer cell lines

To explore the molecular aspect of the alteration of exosome secretion, we first attempted to confirm the data obtained in our transcriptomic and proteomic approaches. In HepG2 cells, we confirmed that transient (Figures sup 2a-b) and stable (Figures 3a-3b) mutated β-catenin depletion increased gene and protein expression of both *SDC4* and *RAB27A*. Due to the lack of efficient antibodies against SDC4, we were unable to analyze SDC4 protein expression by Western-Blot. However, we were able to confirm the increased expression of Syndecan-4 and Rab27a by immunofluorescence (Figures 3c-3d). We then extended these results to other liver cancer cell lines. We analyzed *SDC4*, *RAB27A*, *ARG1* and *AXIN2* basal mRNA expression in five different liver cancer cell lines, each with a different *CTNNB1* mutational status, and identified positive correlations between *SDC4* and *RAB27A* with *ARG1* (β-catenin-negative target) and negative correlations between *SDC4* and *RAB27A* with *AXIN2* (β-catenin-positive target) (Figure sup 2c-d). Upon analyzing the basal expression of Rab27a protein in these five cell lines, we identified a negative correlation between β-catenin and Rab27a expression (Figures sup 2f- 2g). In the two liver cancer cell lines bearing a β-catenin point mutation (Huh6 and SNU398 cells), inducible shRNA strategy with doxycycline reduced β-catenin protein expression by 63% and 49%, respectively (Figures sup 2h-2i). Associated with that reduction in β-catenin protein expression we observed a decrease in *CCND1* and *AXIN2* mRNA expression and an increase in *ARG1* mRNA expression in both cell lines (Figures sup 2j-2k). As in HepG2 cells, β-catenin depletion also increased *SDC4* and *RAB27A* gene expression and Rab27a protein expression in Huh6 and SNU398 cells treated with doxycycline (Figures 3e-h). In order to mimic the β-catenin activation in a non-mutated HCC cell line, we treated Huh7 cells with a GSK3 inhibitor (CHIR99021) limiting β-catenin phosphorylation and degradation. This CHIR99021 treatment increased β-catenin protein expression (2.37-fold) (Figure 3i) and induced its translocation into the nucleus in Huh7 cells (Figure 3j) where concomitant deregulation of β-catenin transcriptional targets was observed (Figure sup 2l). Consistent with our earlier data using HepG2 cells, CHIR99021 treatment decreased *SDC4* and *RAB27A* gene expression, and Rab27a protein level in Huh7 cells (Figures 3k-l). Then, we used TIRF microscopy to analyze exosomal release in living Huh7 cells expressing CD63-pHluorin. We found that CHIR99021 treatment significantly decreased the number of fusion events in Huh7 cells (Figure 3m). All together, these results revealed that β-catenin mutation/activation represses the expression of syndecan-4 and Rab27A, two proteins involved in exosome biogenesis, in association with repression of exosomal secretion.

**Figure 3.**
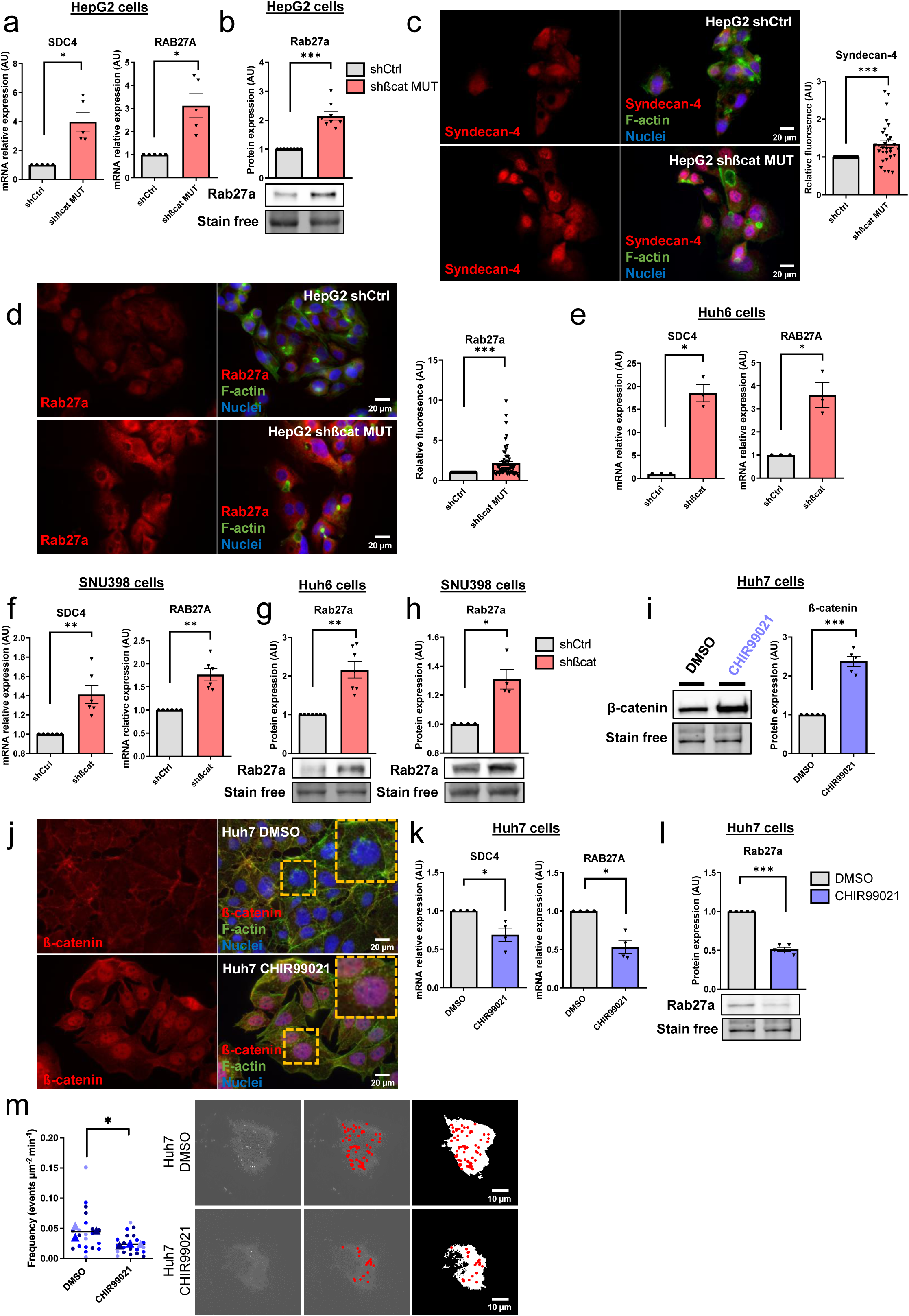
Activated β-catenin represses *SDC4* and *RAB27A* expression in liver cancer cells. (**a-d**) HepG2 cells were treated with doxycycline (DOX) to express either a control shRNA (shCtrl) or a shRNA targeting mutated β-catenin (shβcat MUT). **(a)** Analysis of *SDC4* and *RAB27A* mRNA expression by qRT-PCR. Graphs show the quantification. **(b)** Analysis of Rab27a protein expression by western-blot. Stain free was used as loading control. The graph shows the quantification. **(c-d)** Epifluorescence images of HepG2 shCtrl and shβcat MUT cells stained with Syndecan-4 and Rab27a antibodies (red), Phalloidin (green), Hoechst (blue). Scale bar: 20 µm. Graphs show the quantification of the fluorescence intensity per image and divided per nuclei. Depicted data are representative of three independent experiments, each dot represents one image. **(e)** Analysis of *SDC4* and *RAB27A* mRNA expression in Huh6 cells expressing either a control shRNA (shCtrl) or a shRNA targeting β-catenin (shβcat) treated with DOX. The graph shows the quantification. **(f)** Analysis of *SDC4* and *RAB27A* mRNA expression in SNU398 cells expressing either a control shRNA (shCtrl) or a shRNA targeting β-catenin (shβcat) treated with DOX. The graph shows the quantification. **(g-h)** Analysis of Rab27a protein expression in Huh6 (g) or SNU398 (h) cells expressing either a control shRNA (shCtrl) or a shRNA targeting β-catenin (shβcat). Stain free was used as loading control. The graphs show the quantification. **(i-m)** Huh7 cells treated with DMSO or CHIR99021 (3µM) for 48 hours. **(i)** Analysis of β-catenin expression by western-blot. The graph shows the quantification. **(j)** Epifluorescence images of cells stained with β-catenin antibody (red), Phalloidin (green), Hoechst (blue). Scale bar: 20 µm. **(k)** Analysis of *SDC4* and *RAB27A* mRNA expression by qRT-PCR. The graphs show the quantification. **(l)** Analysis of Rab27a protein expression by western-blot. The graph shows the quantification. **(m)** Quantification of CD63- pHluorin MVB–PM fusion events visualized by live TIRF microscopy. Depicted data are representative of three independent experiments; each dot represents one cell. Images represent the cell mask (white) and red dots corresponding to fusion events. **(a-m)** All graphs show the quantification of at least three independent experiments. Results are expressed as Mean ± SEM, two-tailed Student’s t-test analysis. *P<0.05; **P<0.01; ***P<0.001.

### Mutated β-catenin promotes immune evasion in HCC cells through exosomes

Our next goal was to investigate the possible role of mutated β-catenin in HCC immune escape. We used 3D spheroid models from two liver cancer cell lines (HepG2 and Huh7) and after 24h co-culture of spheroids with Peripheral Blood Mononuclear Cells (PBMC), immune cell infiltration was evaluated by flow cytometry (Figures 4a-4b). First of all, we assessed the capability of PBMC to invade spheroids formed from Huh7 cells treated with CHIR9901. We observed a decrease in PBMC infiltration and an increase in tumor cell survival in Huh7 cells were treated with CHIR9901 compared to the results seen in the control condition (Figure 4c). Of note, we confirmed that CHIR99021 increased β-catenin and decreased Rab27a protein expression in Huh7 spheroids with no impact on the cell viability (Figures sup 3a-3b). Next, we showed that the depletion of mutated β-catenin protein enhanced PBMC infiltration in HepG2 cells associated with a decrease of tumor cell survival (Figure 4d). We verified that mutated β-catenin protein was correctly depleted in HepG2 spheroids, with an increase in Rab27a protein expression and no impact on cell viability (Figures sup 3c-d). Altogether, these results confirm the involvement of mutated β-catenin in immune evasion. We next sought to investigate the role of exosomes in this process. In order to blunt exosome secretion, we used siRNA directed against Rab27a in spheroids formed with HepG2 cells depleted of mutated β-catenin. Rab27a depletion decreased the number of secreted particles without any effect on cell viability in HepG2 shβcat MUT spheroids (Figures sup 3e-g). Furthermore, depletion of Rab27a decreased PBMC infiltration associated with an increase of tumor cell survival (Figure 4e). Taken together, these results strongly suggest a determinant role for exosomes in immune cell infiltration.

**Figure 4.**
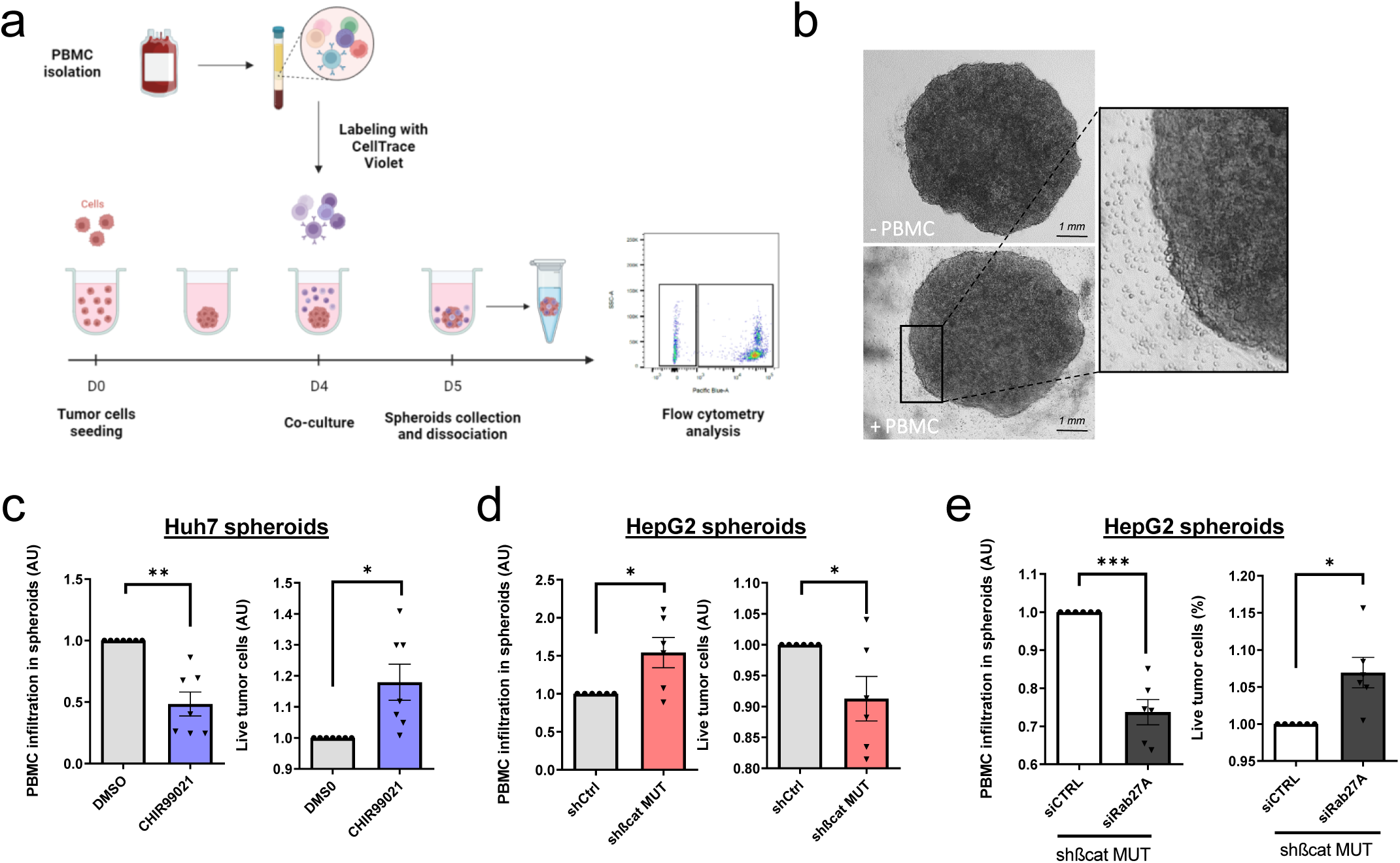
Activated β-catenin represses immune infiltration in liver cancer spheroids through exosomes. (a) Peripheral Blood Mononuclear Cells (PBMC) infiltration analysis protocol. Created with BioRender.com. **(b)** Images of HepG2 spheroids expressing a control shRNA incubated or not with PBMC for 24 hours. **(c)** Analysis of PBMC infiltration and tumor cell survival in Huh7 spheroids treated with DMSO or CHIR99021. **(d)** Analysis of PBMC infiltration and tumor cell survival in HepG2 spheroids expressing control shRNA (shCtrl) or shRNA targeting mutated β- catenin (shβcat MUT). **(e)** Analysis of PBMC infiltration and tumor cell survival in HepG2 spheroids co-expressing shRNA targeting mutated β-catenin (shβcat MUT) and control siRNA (siCtrl) or siRNA targeting Rab27A (siRab27A). (**c-e**) Graphs show the quantification of six independent experiments. Results are expressed as Mean ± SEM, one or two-tailed Student’s t-test analysis. *P<0.05; **P<0.01; ***P<0.001.

### *CTNNB1* mutations are associated with low expression of exosomal biogenesis-associated genes in HCC patient samples

To determine whether markers of exosomal biogenesis are deregulated in HCC human samples, we assessed the expression of *SDC4* and *RAB27A* using public transcriptomic data from two different cohorts (Figure 5a: TGCA data sets from Cbioportal, Figure 5b: Boyault et *al.* article (26)). In both set of data, we showed that *SDC4* and *RAB27A* gene expressions were significantly reduced in *CTNNB1*- mutated HCCs compare to non-mutated tumors (Figures 5a-5b). Furthermore, Pearson’s correlation analysis of these two cohorts revealed that *RAB27A* and *SDC4 gene* expression were positively correlated with the expression of *ARG1* and *PCK1* (β-catenin-negative targets) (27) and negatively correlated with the expression of *GLUL*, *CCND1*, *AXIN2*, *LGR5, FAT1* and *BMP4* (β-catenin-positive targets) (Figures sup 4a-b). We then performed immunohistochemical analysis on 56 human HCC samples with or without *CTNNB1* mutations (Figure sup 4c). We found that tumors strongly positive for glutamine synthetase (a well-known β-catenin-positive target in the liver (28)), were correlated with low levels of Rab27a protein expression (Figure 5c), supporting the link between β-catenin activation and regulation of exosomal secretion through a decrease in Rab27a expression.

**Figure 5.**
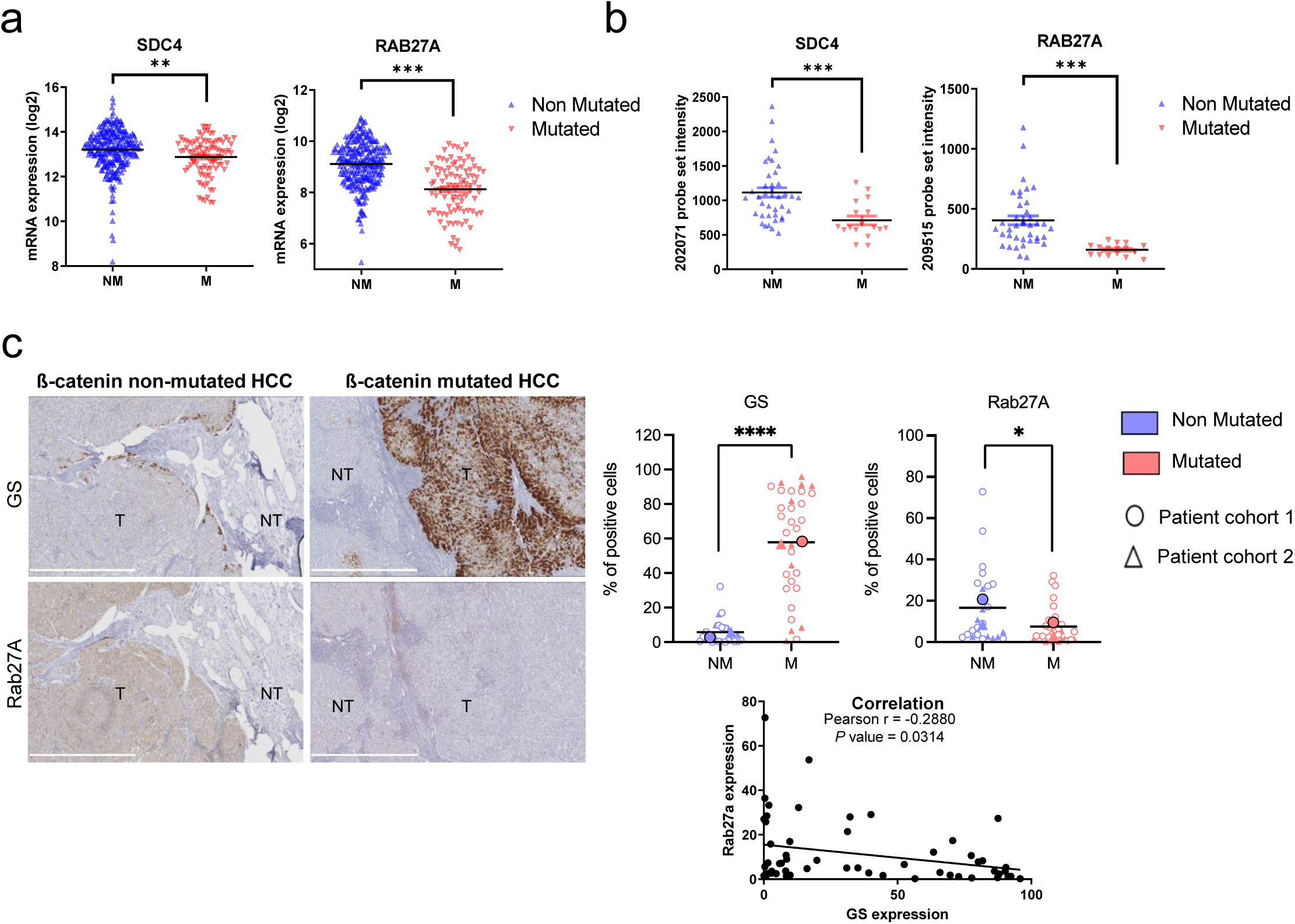
Downregulation of *SDC4* and *RAB27A* in human HCC samples with *CTNNB1* mutations. (a) Analysis of *SDC4* and *RAB27A* mRNA expression in β- catenin mutated (red) and non-mutated (blue) HCC (Cbioportal cohort, n=366). **(b)** Analysis of *SDC4* and *RAB27A* mRNA expression in β-catenin mutated (red) and non-mutated (blue) HCC (Boyault et *al*. cohort, n=56). **(c)** Immunohistochemistry (IHC) analysis of glutamine synthetase (GS) and Rab27a in HCC samples presenting or not β-catenin mutations. Scale bar: 1mm. (T: tumoral, NT: non tumoral). The upper graphs show the quantification of the percentage of cells positive for GS or Rab27A. The analysis was split into two cohorts (circle and triangle). The lower graph shows the Pearson correlation between Rab27a and GS protein expression (n=56). Results are expressed as Mean ± SEM, two-tailed Student’s t-test analysis or Pearson correlation test. *P<0.05; **P<0.01; ***P<0.001; ****P<0.0001.

## DISCUSSION

β-catenin is a main oncogene in tumors (9) and several studies have reported a key role for mutated β-catenin in immune escape in several types of cancer such as HCC, colorectal cancer, melanoma and glioblastoma (6,11,12,15), but its mechanisms of action in this process remain to be elucidated. During the past decades, β-catenin mutated HCC have been classified at the molecular, histopathological and clinical levels as liver cancers with specific features (29–31). More recently, these tumors have been described as cold with low immune infiltrates and resistant to immunotherapy (3–6).

In this study we deciphered the role of mutated β-catenin in cell communication between cancer cells and immune cells. Based on omic results, we hypothesized a role for EV in the control of immune infiltrates in β-catenin-mutated HCC. Following ISEV guidelines (32), we applied various techniques to address the impact of β- catenin activation on EV secretion. Using NTA, we demonstrated that HepG2 cells silenced for mutated β-catenin secrete more particles than do control cells. Our different analyses and results (TIRF with the CD63 exosomal marker, TEM for the EVs shape and size, regulation of Rab27a expression) led us to take a strong interest in small EVs (i.e. exosomes), a subpopulation of EVs. Using real-time visualization of the release of CD63-enriched EVs we demonstrated that mutated β- catenin silencing in HepG2 cells favors small EV secretion. The link between activation of β-catenin and secretion of small EVs was further confirmed in β-catenin non-mutated cells, Huh7 cells, treated with GSK3 inhibitor, suggesting that activation of the Wnt/β-catenin pathway may result in a repression of the EV machinery in tumor cells. This link between β-catenin and EVs was poorly explored in the literature and essentially restricted to the identification of β-catenin as a common EV cargo (33). For example, it was shown that tetraspanins expression reduces the cellular pool of β-catenin by enhancing the exosome-associated export of β-catenin from the cell and that this exosomal discharge of β-catenin downregulates the Wnt signaling pathway (34). More recently, using a genome-wide CRISPR/Cas9-mediated screening to identify genes involved in EV release, Lu et *al*. identified the Wnt signaling pathway in the top hits (35). Indeed, by this approach, a number of Wnt-signaling modulators appeared among the candidates as EV biogenesis regulators, which were further validated in chronic myelogenous leukemia K562 cells. Lu et *al.* also demonstrated that Wnt-mediated GSK3 inactivation, in K562 cells, regulated EV release in a Rab27a dependent manner (35). Herein, we added new findings reporting that this regulation also occurs in solid tumors. Moreover, we demonstrated for the first time the specific involvement of the oncogenic β-catenin in the regulation of exosome biogenesis.

At the molecular level, thanks to global approaches we found that oncogenic β- catenin represses the expression of genes involved in the EV machinery. We indeed found that genes from the Rab (*RAB8B*, *RAB11A*, *RAB11B*, *RAB17, RAB27A*), Vamp (*VAMP2*, *VAMP5*, *VAMP8*) and Vps (*VPS4A*, *VPS4B*, *VPS16*, *VPS28*) families were upregulated upon removal of the oncogenic β-catenin in HepG2 cells. Therefore, several trafficking steps could be impaired along the EV secretion pathway, the MVB biogenesis with the Vps, the transport and docking with the Rab and the final step of membrane fusion with the Vamp (16). Herein, we focused on *RAB27A* and *SDC4* genes, which were the most dysregulated in our transcriptomic analysis. We showed that mutated β-catenin represses their gene expression in liver cancer cell lines. In non-mutated β-catenin HCC cells, activation of β-catenin showed that both gene and protein expression of Rab27a were decreased. We further found a correlation between the presence of *CTNNB1* mutations and the low expression of *RAB27A* and *SDC4* in human HCC patient samples. The way *RAB27A* and *SDC4* gene expression is repressed by β-catenin remains to be explored. We described positive correlation between expression of *SDC4* and *RAB27A* and expression of negative transcriptional targets of β-catenin, as well as a negative correlation with expression of positive targets in tumors. Even if the mechanism by which β-catenin and LEF/TCF transcription factors led to target gene expression activation is well known (21,36,37), it is not the case for the repression process. In melanoma the transcription repressor ATF3 induced by the Wnt/β-catenin pathway has been associated with *CCL4* repression (11). Analysis of *SDC4* and *RAB27A* promoter sequences revealed the presence of consensus sequences of Lef1, Tcf4 and Atf3 binding (data not shown), suggesting that these newly identified target genes could be regulated at the transcriptional level by transcription factors known to associate with β-catenin. In this line, *SDC4* was found to be a negative Tcf4 target gene in HCC mouse models (38).

Following the EV machinery alteration, we further addressed the impact of the β- catenin-mediated repression of EV secretion on immune cell infiltration in tumors. Herein, the functional role of these tumor-derived exosomes in tumor immunity was studied using liver cancer 3D models and PBMC. Using siRNA depletion of Rab27a, we showed that mutated β-catenin promotes a decrease of immune cell infiltration in micro-tumors through a decrease of exosome secretion by . The small GTPase Rab27a has been implicated in several cancer progression mechanisms such as metastasis, cell invasion or migration (39). The oncogenic function of Rab27a is mainly due to its function as regulator of exosome secretion, which modulates cancer cell function and tumor microenvironment. In mammary carcinoma cells, the decrease of exosome secretion by Rab27a inhibition resulted in a diminution of primary tumor growth and lung metastatic dissemination (40). However, shRNA knockdown of Rab27a in a prostate cancer model revealed a dominant immunosuppressive role for EVs during antigen cross presentation (41). These data are inconsistent with our own, meaning so it seems that Rab27a-induced exosomal secretion could be a “double-edged sword” in tumor progression. This could be due to the content of EVs. In fact, EVs are known to contain heterogeneous cargos that can play a critical role in immunomodulation (42). For example, the main immunosuppressive cytokine TGF-β expressed by tumoral-derived EVs has been identified in Natural Killer cell function suppression (43,44). EVs released by tumoral cells can also carry several miRNAs mainly known to be involved in immune suppression (42). MiR-103a and Let-7a in tumor EVs promote M2-like polarization in lung and melanoma cancers (45,46). Interestingly, immunomodulatory proteins such as CSF-1, CCL2, FTH, FTL, and TGF-β chemokines have been identified in tumor exosomes (46). The content of β-catenin-regulated EVs remains to be explored to fully understand their function in the immunomodulation of the tumor microenvironment.

In conclusion, this study is the first description of the oncogenic β-catenin as a key regulator of the exosome machinery. Moreover, it gave new insights regarding the regulation of tumor-derived exosome production by the β-catenin pathway and their involvement in the immune escape observed in β-catenin mutated HCC. Given that β-catenin is also mutated in melanoma, endometrial and colorectal cancers, these new findings may have a broader significance. Our data indicate that exosomes could be a promising biomarker and that restoring the tumor exosomal secretion of β- catenin-mutated tumors may increase the communication between tumor and immune cells, and as a result, turn cold tumors hot.

## Supporting information

Video TIRF HepG2 shCTRL

Video TIRF HepG2 shMUT

Video TIRF Huh7 DMSO

Video TIRF Huh7 CHIR99021

## Acknowledgments

We thank the FACSility, OneCell, Histopathologie and Oncoprot facility platforms (TBMCore, UMS005, Bordeaux) for the help with flow cytometry, qPCR, immunohistochemistry and mass spectrometry experiments. We acknowledge Cyril Dourthe for his help in the bioinformatic analysis. We thank Silvia Sposini (IINS, Bordeaux) for her help in the TIRF microscopy experiments. We thank Drs. Sara Basbous and Benjamin Bonnard (BRIC, Bordeaux) for helpful discussions on the project. We are grateful to Drs. Jean-Christophe Delpech, Liam Barry-Carroll (NutriNeuro Laboratory, Bordeaux) and Alexandre Favereaux (IINS, Bordeaux) for insightful discussions about EVs. We acknowledge the Bordeaux Imaging Center (Bordeaux, France) and the France BioImaging infrastructure supported by the French National Research Agency (ANR-10-INSB-04, “Investments for the future”). We thank Professor William A. Thomas (Colby-Sawyer College, New Hampshire) for proof-reading of the manuscript. CD is supported by a PhD fellowship from both SIRIC BRIO and Région Nouvelle-Aquitaine. JV is supported by PhD scholarships from the French Ministry of Research (MENESR). This research was funded by La Fondation pour la Recherche Médicale to VM (équipe FRM 2018, grant number DEQ20180839586), La Ligue contre le Cancer (comité des Charentes et comité de la Gironde) to CB and l’AFEF (Association Française de l’Etude du Foie) to CB. The authors declare no competing financial interests.

## Abbreviations

BC: bile canaliculi
DOX: doxycycline
EV: extracellular vesicle
FC: fold change
GS: glutamine synthetase
HCC: hepatocellular carcinoma
IHC: immunohistochemistry
KD: knockdown
MVB: multivesicular body
NTA: nanoparticle tracking analysis
PBMC: Peripheral Blood Mononuclear Cell
PM: plasma membrane
TEM: transmission electron microscopy
TIRF: total internal reflection fluorescence.

## Author contributions

Study design: CB, VM. Generation of experimental data: CD, JV, AB, IM, MP, NDS, CB. Analysis and interpretation of data: CD, JV, AB, AAR, JWD, PB, DP, MV, BB. Writing of the manuscript: CD, JV, CB, VM. Supervision of the project: CB, VM.

## MATERIALS AND METHODS

### Cell culture

Human HepG2 and Huh7 cell lines were grown in 4.5 g/L glucose Dulbecco’s modified Eagle’s Medium (DMEM, Gibco) supplemented with 10 % FBS (Fetal Bovine Serum, Sigma). Human Huh6 cells were grown in 1 g/L glucose DMEM, (Gibco) supplemented with 10 % FBS. Human SNU398 cells were grown in Roswell Park Memorial Institute medium (RPMI 1640, Gibco) supplemented with 10 % FBS. Heat-inactivated FBS was used (30 min, 56°C). All cell lines were cultured in a humidified atmosphere containing 5% CO_2_ and 37°C and mycoplasma contamination was checked regularly by PCR. All cell lines, except Huh6, were purchased from American Type Culture Collection (ATCC). The Huh6 cell line was generously provided by C. Perret (Paris, France).

### Transfection

DNA transfections were performed using the Lipofectamine 3000 transfection Reagent according to the manufacturer’s instructions (Invitrogen). The following plasmid was used: pCMV-Sport6-CD36-pHluorin (Plasmid#130901, Addgene). siRNA oligos were transfected into cells using Lipofectamine RNAiMax Reagent according to the manufacturer’s protocol (Invitrogen). A reverse transfection was performed on day 1, a second forward transfection on day 2 and experiments were conducted on day 5. siRNA β-catenin MUT (Eurofins Genomics) was previously reported (21), siRNA RAB27A (MISSION EHU091501, Sigma) targets human RAB27A. The AllStars negative-control siRNA from Qiagen was used as control siRNA.

### Lentiviral infection

HEK293T cells were seeded (2.5x10^6^) on 10 cm plate coated with poly-L-lysine (Sigma) to obtain a confluence of 50-70% at the time of the infection. The next day, 600 µL of Opti-MEM (Gibco), 22 µL of Mirus LT1 transfection reagent (Mirus) and 4 µg of lentiviral plasmids of packaging vector (pPAX, pSD11) and lentiviral plasmid of interest were added on cells and incubated for 48h to allow the production of viruses. Then supernatant was harvested, centrifuged at 2500 rpm for 3 min to remove dead cells and debris, filtered and used directly to infect cells or stored at -80°C. After infection with lentiviral particles, cells were selected with puromycin treatment (2 µg/mL) for 1 week.

### shRNAs

The Tet-pLKO-puro lentivirus vector (plasmid<u>#21915)</u> and the control shRNA lentivirus vector (pLKO-Tet-On-shRNA-Control, plasmid<u>#398398)</u> were purchased from Addgene (Watertown, USA). The construction of the two shRNA lentivirus vectors targeting the human β-catenin was performed following the Tet-pLKO Manual given by Addgene (plasmid<u>#21915)</u>. The shRNA sequences used were the same as previously described (16): the 5’-ACCAGTTGTGGTTAAGCTCTT-3 sequence to target the human β-catenin in Huh-6 and SNU398 cells, and the 5’- TGTTAGTCACAACTATCAAGA-3’ sequence to target specifically the mutated form of β-catenin in HepG2 cells. shRNAs were induced with doxycycline (1µg/mL) in Huh6 cells for 5 days (2 treatments) and in HepG2 and SNU398 cells for 7 days (3 treatments).

### Drug treatment

Huh7 cells were treated with the GSK3 inhibitor CHIR99021 (3 µM, Sigma) for 48h before cell analysis. DMSO was used as control.

### Spheroid formation

Human Huh7 cells treated with either DMSO or CHIR99021 (3 µM; Sigma) for 48h were seeded (20,000 cells per well in 100µL) in non-adherent conditions (ultra-low attachment 96 wells plate, Costar) in filtered 4.5 g/L glucose Dulbecco’s modified Eagle’s Medium (DMEM, Gibco) supplemented with 10 % exosome free FBS. Human HepG2 cell line was transfected with either control shRNA or shRNA directed against mutated β-catenin (shBcat MUT) and induced with doxycycline (1 µg/mL, 3 rounds in one week). HepG2 shβcat MUT were also transfected with siRNA targeting Rab27a the day before the seeding for the reverse transfection (day 0) and the day of the seeding for the second forward transfection (day 1). 10,000 HepG2 cells were seeded per well in non-adherent conditions (ultra-low attachment 96 wells plate, Costar) in 100 µL filtrated 4.5 g/L glucose Dulbecco’s modified Eagle’s Medium (DMEM, Gibco) supplemented with 10 % exosome free FBS for 96 hours. The induction of shRNA was maintained by adding doxycycline in each well (1µg/mL in 50µL) the day of the seeding and 48 hours after.

### Peripheral Blood Mononuclear Cell (PBMC) infiltration

After 96 hours of formation, Huh7 and HepG2 spheroids were co-cultured with CellTrace Violet (Life Technologies) labeled Peripheral Blood Mononuclear Cells (PBMC). Briefly, PBMC were isolated from donor blood (EFS Bordeaux) using a Ficoll (Eurobio Scientific) density gradient centrifugation. The obtained cells were then centrifuged (10 min, 1500 rpm) and washed several times in PBS 1X. The remaining red blood cells were lysed by incubation in an ACK lysing buffer (Gibco), and the platelets were eliminated by centrifugation (10 min, 900 rpm). PBMC were then stained with Cell Trace Violet (5 μM, 20 min, 37°C, 10^6^ cells/mL) and added to the wells (ratio tumor cells per spheroid:PBMC = 1:5, in 50 µL) for a 24 hours co-culture. After 24 hours, spheroids were collected, washed twice with PSB 1X and dissociated with trypsin 0.25%-EDTA (10 min, 37°C, Gibco). Cells were then pelleted by centrifugation, resuspended in 200 µL of MACS buffer (PBS 1X Dutscher, 0.5% FBS Eurobio Scientific, 0.4% EDTA 0.5 mM Euromedex) and labelled with Propidium Iodide (1/500 dilution, Sigma-Aldrich). The proportion of immune cells infiltrated in spheroids and tumor cells survival was then analyzed by flow cytometry (CantoII cytometer, BD Biosciences, Le Pont de Claix, France) and data analysis was performed with the FlowJo software (version 10.8.1).

### qRT-PCR analysis

Total RNA was extracted from cells using the kit NucleoSpin RNA (Macherey-Nagel). RNA was then reverse transcribed using material from ThermoFisher Scientific. qPCR was performed using the SYBR Green SuperMix (Quanta) using a C1000 Real-Time System (Bio-Rad). Data were normalized using the r18S gene as endogenous control and fold change was calculated using the comparative Ct method (-ddCt). All primers used are listed in table 1.

**Table 1.**
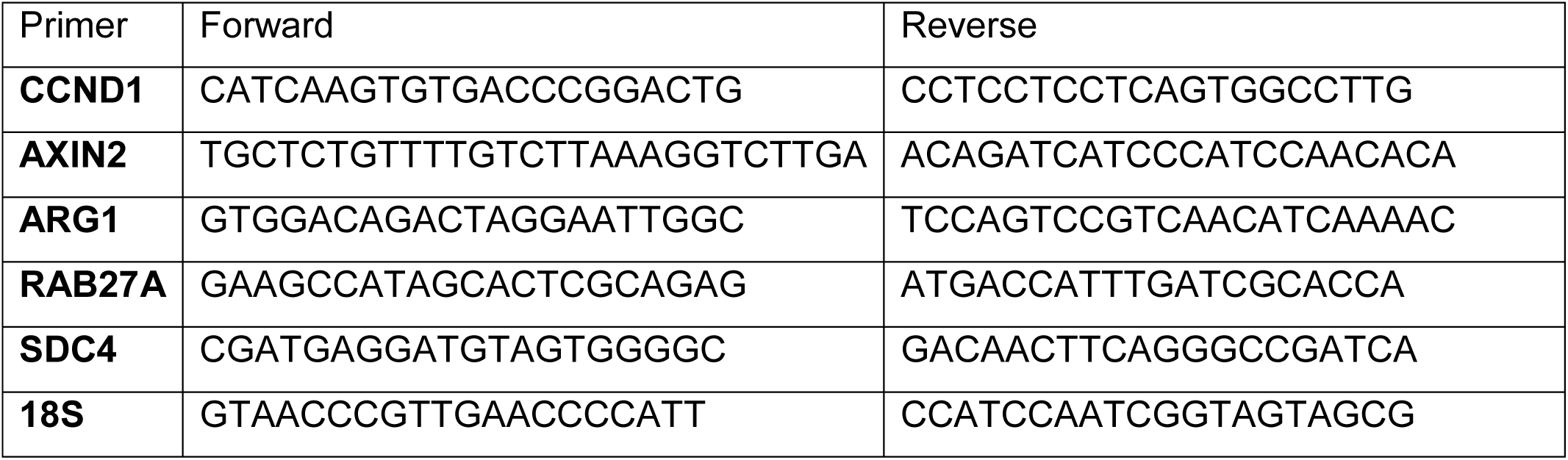
qPCR primers.

### Western blot

Total proteins were extracted from cells or EVs using RIPA lysis buffer (0.1% SDS, 1% NP40, 0.15 M NaCl, 1% sodium deoxycholate, 25 mM Tris HCl pH 7.4) supplemented with protease and phosphatase inhibitors (Roche). Proteins were then denaturated in Laemmli buffer (Bio-Rad) at 95°C for 5 min. 40 µg of protein extract was loaded on 10% polyacrylamide gels (TGX Stain-Free FastCast, Bio-Rad). Membranes were blocked with 5% BSA-TBST (5% BSA Sigma, TBS 1x Euromedex, 0,1% Tween20 Sigma) for 30 min and incubated with primary antibodies (diluted in 5% BSA-TBST) overnight at 4°C. The following primary antibodies were used: β- catenin (1:2,000, mouse, 610154, BD Biosciences), Cyclin D1 (1:1,000, mouse, sc- 20044, Santa Cruz), Rab27a (1:1,000, rabbit, 69295, Cell signaling), CD63 (1:500, rabbit, SAB430160, Sigma). Membranes were then incubated with secondary antibodies (1:5,000 diluted in 5% BSA-TBST) for 30 min. The following secondary antibodies were used: IRDye 680CW conjugated goat anti-rabbit IgG (H&L) (LI-COR), IRDye 800CW conjugated goat anti-mouse IgG (H&L) (LI-COR). Acquisitions were performed using the ChemiDoc Imaging System (Bio-Rad). Intensities were measured using the ImageLab software (Bio-Rad).

### EVs isolation and characterization

Cells (2 million) were cultured in medium (7 mL) supplemented with EV-depleted FBS (obtained by ultracentrifugation for 16 h at 120,000 g, using an Optima XPN-80 centrifuge with a 45.Ti rotor). EVs were isolated 72 hours after by differential centrifugation at 4°C: 5 min at 300 g, 5 min at 2000 g, 40 min at 18,000 g and 120 min at 120,000 g. EV pellets were washed in PBS, centrifuged again at 120,000 g for 120 min, resuspended in PBS (or RIPA lysis buffer) and stored at -80°C.

### Nanoparticle tracking analysis (NTA)

Number and size of particles were detected using the NS300 instrument (Malvern Panalytical Ltd., Malvern, UK) equipped with a 488 nm laser and a high-sensitivity scientific CMOS camera. Particles are tracked and sized based on Brownian motion and diffusion coefficient. Particles were diluted in PBS to obtain a concentration within the recommended range (30-120 particles/frame). Five 60s videos were acquired for each sample with the following condition: cell temperature 25 °C, syringe speed—22 µL/s, camera level 14. Videos were subsequently analyzed with NanoSight Software NTA3.3.301 (Malvern Panalytical Ltd., Malvern, UK), which identified and tracked the center of each particle under Brownian motion to measure the average distance the particles moved on a frame-by-frame basis.

### Electron microscopy

#### Cells

Cells were fixed in EM fixative solution (1.6% Ga in PB 0.1M pH 7.4) for 1h at room temperature and then centrifuged. Cell pellets were embedded in 1% agarose. Samples were washed three times with PB 0.1M and postfixed in 1.5% potassium ferrocyanide, 2% aqueous osmium tetroxide solution in PB 0.1M for 1h at room temperature. Before and between each of the following incubations, samples were washed three times with distilled water. Samples were first incubated in fresh thiocarboxyhydrazide (TCH) solution for 20 min at room temperature. In a second postfixation step, a 2% osmium tetroxide solution was added on samples for 30 min at room temperature. Samples were then serially dehydrated in ethanol (35%, 35%, 50%, 70%, 80%, 90%, 100%, 100%, 100%; 2 min each), incubated in a solution of 50% ethanol and 50% Epon resin for 2 hours and incubated in 100% Epon resin overnight at room temperature. Epon resin was then renewed for another 4 hours incubation. At the end, samples were embedded in a pure resin and cured by incubation at 60°C for 48 hours. Ultramicrotomy was done with a Diatome 35° Diamond knife and a Leica EM-UC7 ultracut. Sections (70 nm thick) were collected on 200mesh copper grid. Images were acquired using a transmission electron microscope (Hitachi H7650) at 80Kv equipped with an Orius Camera (Gatan) managed by Digital Micrograph software. The quantification of the number of MVBs was performed by counting visible MVBs and then dividing per the cell surface using ImageJ software. The quantification of the MVB diameter was performed using ImageJ software.

#### EVs

Cell supernatant fluid was concentrated using an Amicon Ultra-15 centrifugal filter from 6 mL to 500 µL at 4000 g during 30 min (100k, Merk Millipore). EVs were isolated using the IZON qEV original 35 nm size exclusion column (Izon science) according to the manufacturer’s instructions. EVs were then concentrated as mentioned previously and resuspended in an equal volume of 4% PFA in PBS (2% PFA final). 20 µL drops of the resuspended exosomes were then absorbed for 20 min at RT on hydrophilic carbon coated grids. After PBS rinses, grids were placed on drops of 1% glutaraldehyde in PBS for 5 min. Several washes were done with distilled water, and grids were then transferred on a filtered drop of pH7 uranyl-oxalate solution, for 5 min in the dark. Grids were then transferred in drops of filtered 4% aqueous uranyl acetate/2,3M methylcellulose (1V for 9V) on a petri dish covered with parafilm on ice for 10 min in the dark. Grids were then removed, one at time, with a stainless-steel loop, and excess fluid were blotted by gently touching the edge of the loop with a Whatman No. 1 filter paper. Grids were air-dried while still on the loop, torn off and stored in a grid storage box. Observations were done with a TEM Hitachi H7650 at 80kV equipped with a Gatan Orius camera.

### Label-free quantitative proteomics

Three independent biological replicates were performed on total protein extracts from human HepG2 cells transfected with control siRNA or β-catenin mutated siRNA. 10 μg of proteins were loaded on a 10% acrylamide SDS-PAGE gel and proteins were visualized by Colloidal Blue staining. Migration was stopped when samples had just entered the resolving gel and the unresolved region of the gel was cut into only one segment. The steps of sample preparation and protein digestion by the trypsin were performed as previously described (44). NanoLC-MS/MS analysis were performed using an Ultimate 3000 RSLC Nano-UPHLC system (Thermo Scientific, USA) coupled to a nanospray Orbitrap Fusion™ Lumos™ Tribrid™ Mass Spectrometer (Thermo Fisher Scientific, California, USA). Each peptide extract was loaded on a 300 μm ID x 5 mm PepMap C18 precolumn (Thermo Scientific, USA) at a flow rate of 10 μL/min. After a 3 min desalting step, peptides were separated on a 50 cm EasySpray column (75 μm ID, 2 μm C18 beads, 100 Å pore size, ES903, Thermo Fisher Scientific) with a 4-40% linear gradient of solvent B (0.1% formic acid in 80% ACN) in 57 min. The separation flow rate was set at 300 nL/min. The mass spectrometer operated in positive ion mode at a 2.0 kV needle voltage. Data were acquired using Xcalibur 4.4 software in a data-dependent mode. MS scans (m/z 375- 1500) were recorded at a resolution of R = 120000 (@ m/z 200), a standard AGC target and an injection time in automatic mode, followed by a top speed duty cycle of up to 3 seconds for MS/MS acquisition. Precursor ions (2 to 7 charge states) were isolated in the quadrupole with a mass window of 1.6 Th and fragmented with HCD@28% normalized collision energy. MS/MS data was acquired in the ion trap with rapid scan mode, a 20% normalized AGC target and a maximum injection time in dynamic mode. Selected precursors were excluded for 60 seconds. Protein identification was done in Proteome Discoverer 2.5. Mascot 2.5 algorithm was used for protein identification in batch mode by searching against a Uniprot Homo sapiens database (75 793 entries, release September 3, 2020). Two missed enzyme cleavages were allowed for the trypsin. Mass tolerances in MS and MS/MS were set to 10 ppm and 0.6 Da. Oxidation (M) and acetylation (K) were searched as dynamic modifications and carbamidomethylation (C) as static modification. Raw LC-MS/MS data were imported in Proline Studio for feature detection, alignment, and quantification (45). Protein identification was accepted only with at least 2 specific peptides with a pretty rank=1 and with a protein FDR value less than 1.0% calculated using the “decoy” option in Mascot. Label-free quantification of MS1 level by extracted ion chromatograms (XIC) was carried out with parameters indicated previously. The normalization was carried out on median of ratios. The inference of missing values was applied with 5% of the background noise. The mass spectrometry proteomics data have been deposited to the ProteomeXchange Consortium via the PRIDE (46) partner repository with the dataset identifier : PXD043841.

### Immunofluorescence

Cultured cells were fixed with 4% PFA for 10 min, permeabilized with Triton-X100 0.1% (Sigma) for 10 min and incubated in 4% BSA-PBS for 15 min to prevent non-specific binding. Cells were then incubated at room temperature with primary antibodies (diluted in 4% BSA-PBS) for 45 min and with secondary antibodies (1:200 diluted in 4% BSA-PBS) for 30 min. The following primary antibodies were used: β- catenin (1:400, mouse, 610154, BD Biosciences), Rab27a (1:800, rabbit, 69295, Cell signaling), SDC4 (1:200, rabbit, 12236, Cell signaling). The following secondary antibodies were used: 547H donkey anti-mouse IgG (H&L) (FP-SB4110-T, Interchim) and 547H donkey anti-rabbit IgG (H&L) (FP-SB5110, Interchim). Nuclei were stained with Hoechst (1:1,000, 34580, Sigma) and actin was stained with 488 phalloidin (1:200, FP-YE5180, Interchim). Coverslips were mounted on microscope slides using Fluoromount-G mounting media (SouthernBiotech) and were imaged under epifluorescence microscope (Zeiss) using the 63X oil immersion objective and SP5 confocal microscope (Leica). Images were analyzed using ImageJ software.

### Immunohistochemistry

#### Human samples

The total number of human HCC samples is 56 and has been provided by 2 cohorts obtained at different times (Figure sup 4c). Experiments for each cohort have been performed by two different experimenters (cohort 1: NDS, cohort 2: IM) and using two different batches of Rab27a antibody.

#### Experiment

Paraffin-embedded human HCC samples were cut into 3.5-µm thick tissue sections and processed for immunohistochemistry with the EnVision FLEX kit material (K800021-2, Agilent Dako) according to the manufacturer’s instructions. Briefly, tissue sections were put on slides (TOM-1190, Matsunami) and placed on automatic staining racks at room temperature to be deparaffinised. Slides were immersed in a 75°C pre-warmed Target Retrieval Solution Low pH (50x) (K8005) in the Dako PTlink tank. The tank was then heated up to 95°C, slides were incubated for 20 min and allowed to cool to 75°C. Afterwards, tumor sections were soaked for 5 to 10 min in Dako Wash Buffer. Slides were placed on the automatic staining racks of the Dako Autostainer and peroxidase blocking reagent was added for 10 min. Sections were rinsed with Dako Wash Buffer and antibodies were applied for 45 min according to the following dilutions (Dako Antibody Diluent): Glutamine synthetase (1:400, pH6, mouse, 610517, BD Biosciences), Rab27a (1:100, pH6, rabbit, 69295, Cell Signalling), SDC4 (1:200, pH6, P11820-1-A, Proteintech). Two solutions were then applied to the slides with washing with Dako Wash Buffer before and between each application: HorseRadish Peroxydase for 20 minutes and substrate working solution. Finally, slides were rinsed with water and hematoxyline counterstaining was performed before mounting on coverslips (Eukitt classic mounting medium).

### Total internal reflection fluorescence (TIRF) imaging and analysis

Coverslips were placed in an imaging chamber, perfused at 37°C with Hepes Buffer Saline (HBS) solution (135 mM NaCl, 5 mM KCl, 0.4 mM MgCl_2_, 1.8 mM CaCl_2_, 1 mM D-glucose, and 20 mM HEPES) and were adjusted to pH 7.4 and to 305 mOsm. Imaging was performed with an Olympus IX83 inverted microscope equipped for (TIRF) microscopy with a 100×, 1.49 NA objective (UAPON100XOTIRF), a laser source (Cobolt Laser 06-DPL 473 nm, 100 mW), and an ILas2 illuminator (Gataca Systems) with a penetration depth set to 100 nm. Emitted fluorescence was filtered with a dichroic mirror (R405/488/561/635) and an emission filter (ET525/50m, Chroma Technology) and recorded by an electron-multiplying charge-coupled device (EMCCD) camera (QuantEM 512C, Teledyne Photometrics). Movies were acquired for 2 min at 10 Hz for exocytosis. To achieve good signal/noise ratio required for event detection and further analysis, fluorescence was bleached by high laser power illumination prior to acquisition of the full movie. MetaMorph 7.8 software was used for all acquisitions. Semi-automatic detection of exocytic events and their quantification were conducted using custom-made MATLAB scripts previously described (47–49).

### Bioinformatic analysis

Public transcriptomic data of 366 HCC patient samples were provided from the Cancer Genome Atlas Research Network, downloaded from the cBioPortal site and divided in two groups according to the presence or not of *CTNNB1* hotspot mutations (respectively 94 and 272 samples per group) by Genome data. Public transcriptomic data of 56 HCC patient samples were provided from the *Boyault et al.* (26) article and divided in two groups according to the presence or not of *CTNNB1* mutations (respectively 17 and 39 samples per group).

### Statistical analysis

Data are expressed as mean ± SEM and are representative of at least three experiments. Statistical tests were carried out using GraphPad Prism software version 8.0.2 (GraphPad Software, San Diego, CA, USA). Statistical significance (*p* <0.05 or less) was determined using Student’s *t-test* or analysis of variance (ANOVA). Values of *p* are indicated as follows: ∗*p* <0.05; ∗∗*p* <0.01; ∗∗∗*p* <0.001; \*\*\*\**p*<0.0001; ns, non-significant.

## Supplemental data

**Supplemental figure 1. HepG2 model validation (a-d)** HepG2 cells were treated with doxycycline (DOX) to express either a control shRNA (shCtrl) or a shRNA targeting mutated β-catenin (shβcat MUT). **(a)** Analysis of *CCND1*, *AXIN2* and *ARG1* mRNA expression by RT-qPCR. The graph shows the quantification of six independent experiments. **(b)** Cells were stained using fluorescent phalloidin and imaged by confocal microscopy. Images are showing bile canaliculi (BC). Scale bar: 5 µm. **(c)** Quantification of BC area, perimeter and percentage of cells forming BC. Depicted data are representative of three independent experiments with twenty BC per experiment, each dot represents one BC. **(d)** Nanoparticle tracking analysis of supernatant from cells. The graphs show the quantification of the number of particles (left-hand) and of the particle size (right-hand) of seven independent experiments. **(e)** CD63-pHluorin MVB–PM fusion events visualized by live TIRF microscopy of HepG2 shβcat MUT cells. Scale bar: 1 µm. **(f)** Transmission electron microscopy images of HepG2-derived EVs by wide-field (yellow arrowheads). Scale bar: 500nm. Results are expressed as Mean ± SEM, two-tailed Student’s t-test analysis. *P<0.05; **P<0.01; ***P<0.001; ns, non-significant.

**Supplemental figure 2. Correlation of expression of *SDC4* and *RAB27A* with β- catenin targets in liver cancer models and Huh6, SNU398, Huh7 model validation** (a-b) Analysis of *SDC4* and *RAB27A* mRNA expressions (a) and of β- catenin and Rab27a protein expressions (b) in HepG2 cells transfected with control (siCtrl) or mutated β-catenin targeting (siβcat MUT) siRNA by qRT-PCR (a) and Western-blot (b). The graphs show the quantification of at least four independent experiments. **(c)** Analysis of *SDC4* and *RAB27A* basal mRNA expression in HepG2, SNU398, Huh6, Huh7 and Hep3B cell lines by qRT-PCR. The graphs show the quantification of five independent experiments. **(d)** Pearson’s correlation analysis of *SDC4* and *RAB27A* with *ARG1* basal mRNA expression in HepG2, SNU398, Huh6, Huh7 and Hep3B cell lines. **(e)** Pearson’s correlation analysis of *SDC4* and *RAB27A* with *AXIN2* basal mRNA expression in HepG2, SNU398, Huh6, Huh7 and Hep3B cell lines. **(f-g)** Basal expression of β-catenin and Rab27a in liver cancer cell lines mutated (HepG2, SNU398, Huh6) or not (Huh7, Hep3B) for β-catenin analysed by western-blot. Stain free was used as loading control. Graphs show the quantification of three independent experiments. On the right-hand graph, protein expression was subjected to Pearson’s correlation coefficient analysis. **(h-i)** Analysis of β-catenin protein expression in Huh6 (h) and SNU398 (i) cells expressing either a control shRNA (shCtrl) or a shRNA targeting β-catenin (shβcat) treated with doxycycline (DOX). The graphs show the quantification of nine (h) and three (i) independent experiments. **(j-k)** Analysis of *CCND1*, *AXIN2* and *ARG1* mRNA expression in Huh6 (j) and SNU398 (k) shCtrl and shβcat cells treated with DOX. The graph shows the quantification of three independent experiments. **(l)** Analysis of *AXIN2* and *ARG1* mRNA expression in Huh7 cells treated with DMSO or CHIR99021 (3µM). The graph shows the quantification of four independent experiments. Results are expressed as Mean ± SEM, one or two-tailed Student’s t-test analysis. *P<0.05; **P<0.01; ***P<0.001.

**Supplemental figure 3. HepG2 and Huh7 3D model validation. (a)** β-catenin and Rab27a expressions were analyzed by western-blot in Huh7 spheroids treated with DMSO or CHIR99021 (3µM). **(b)** Cell viability of Huh7 spheroids (DMSO and CHIR99021). **(c)** HepG2 cells were treated with doxycycline (DOX) to express either a control shRNA (shCtrl) or a shRNA targeting mutated β-catenin (shβcat MUT) and grown in spheroids. β-catenin and Rab27a expressions were analyzed by western- blot. **(d)** Cell viability of HepG2 shCtrl and shβcat MUT spheroids. **(e)** Analysis of Rab27a expression in HepG2 spheroids co-expressing shβcat MUT and control siRNA (siCtrl) or siRNA targeting Rab27A (siRab27A). **(f)** Cell viability of HepG2 shβcat MUT spheroids treated with siCtrl or siRab27a. **(g)** Nanoparticle tracking analysis of supernatant from HepG2 shβcat MUT spheroids treated with siCtrl or siRab27a. All graphs show the quantification of at least three independent experiments. Results are expressed as Mean ± SEM, one or two-tailed Student’s t- test analysis. *P<0.05; **P<0.01; ***P<0.001; ns., non-significant.

**Supplemental figure 4. Correlation of expression of *SDC4* and *RAB27A* with various β-catenin targets in human HCC samples with *CTNNB*1 mutations. (a)** Analysis of *SDC4* and *RAB27A* mRNA expression in β-catenin-mutated (red) and non-mutated (blue) HCC (Cbioportal cohort n=366). **(b)** Analysis of *SDC4* and *RAB27A* mRNA expression in β-catenin mutated (red) and non-mutated (blue) HCC (Boyault et *al*. cohort n=56). Results are expressed as Pearson correlation test. *P<0.05; **P<0.01; ***P<0.001; ****P<0.0001; ns, non-significant. **(c)** Description of patient cohorts used for the immunohistochemical analysis.

## Videos

HepG2-shCtrl (video 1), HepG2-shβCat MUT (video 2) and Huh7 treated with DMSO (video 3) or CHIR99021 (video 4) cells were transfected with CD63-pHluorin corresponding to Fig. 2c and 3m. Cells were recorded under TIRF illumination on a microscope (IX83; Olympus) for 2 min. Videos show exocytic events (suddenly appearing fluorescent dots) corresponding to MVB-PM fusion.

## References

1. Sung H, Ferlay J, Siegel RL, Laversanne M, Soerjomataram I, Jemal A, et al. Global Cancer Statistics 2020: GLOBOCAN Estimates of Incidence and Mortality Worldwide for 36 Cancers in 185 Countries. CA Cancer J Clin. 2021;71(3):209–49.

2. Finn RS, Qin S, Ikeda M, Galle PR, Ducreux M, Kim TY, et al. Atezolizumab plus Bevacizumab in Unresectable Hepatocellular Carcinoma. N Engl J Med. 2020;382(20):1894–905.

3. Akasu M, Shimada S, Kabashima A, Akiyama Y, Shimokawa M, Akahoshi K, et al. Intrinsic activation of β-catenin signaling by CRISPR/Cas9-mediated exon skipping contributes to immune evasion in hepatocellular carcinoma. Sci Rep. 2021;11(1):16732.

4. Pinyol R, Sia D, Llovet JM. Immune Exclusion-Wnt/CTNNB1 Class Predicts Resistance to Immunotherapies in HCC. Clin Cancer Res Off J Am Assoc Cancer Res. 2019;25(7):2021–3.

5. Ruiz de Galarreta M, Bresnahan E, Molina-Sánchez P, Lindblad KE, Maier B, Sia D, et al. β-Catenin Activation Promotes Immune Escape and Resistance to Anti– PD-1 Therapy in Hepatocellular Carcinoma. Cancer Discov. 2019;9(8):1124–41.

6. Sia D, Jiao Y, Martinez-Quetglas I, Kuchuk O, Villacorta-Martin C, Castro de Moura M, et al. Identification of an Immune-specific Class of Hepatocellular Carcinoma, Based on Molecular Features. Gastroenterology. 2017;153(3):812–26.

7. Coste A de L, Romagnolo B, Billuart P, Renard CA, Buendia MA, Soubrane O, et al. Somatic mutations of the β-catenin gene are frequent in mouse and human hepatocellular carcinomas. Proc Natl Acad Sci U S A. 1998;95(15):8847–51.

8. Rebouissou S, Franconi A, Calderaro J, Letouzé E, Imbeaud S, Pilati C, et al. Genotype-phenotype correlation of CTNNB1 mutations reveals different β-catenin activity associated with liver tumor progression. Hepatology. 2016;64(6):2047.

9. Kim S, Jeong S. Mutation Hotspots in the β-Catenin Gene: Lessons from the Human Cancer Genome Databases. Mol Cells. 2019;42(1):8–16.

10. Yaguchi T, Goto Y, Kido K, Mochimaru H, Sakurai T, Tsukamoto N, et al. Immune suppression and resistance mediated by constitutive activation of Wnt/β- catenin signaling in human melanoma cells. J Immunol Baltim Md 1950. 2012;189(5):2110-7.

11. Spranger S, Bao R, Gajewski TF. Melanoma-intrinsic β-catenin signalling prevents anti-tumour immunity. Nature. 2015;523(7559):231-5.

12. Du L, Lee JH, Jiang H, Wang C, Wang S, Zheng Z, et al. β-Catenin induces transcriptional expression of PD-L1 to promote glioblastoma immune evasion. J Exp Med. 2020;217(11):e20191115.

13. Cen B, Wei J, Wang D, Xiong Y, Shay JW, DuBois RN. Mutant APC promotes tumor immune evasion via PD-L1 in colorectal cancer. Oncogene. 2021;40(41):5984–92.

14. Rotman J, Heeren AM, Gassama AA, Lougheed SM, Pocorni N, Stam AGM, et al. Adenocarcinoma of the Uterine Cervix Shows Impaired Recruitment of cDC1 and CD8+ T Cells and Elevated β-Catenin Activation Compared with Squamous Cell Carcinoma. Clin Cancer Res Off J Am Assoc Cancer Res. 2020;26(14):3791–802.

15. Wang C, Yan J, Yin P, Gui L, Ji L, Ma B, et al. β-Catenin inhibition shapes tumor immunity and synergizes with immunotherapy in colorectal cancer. Oncoimmunology. 2020;9(1):1809947.

16. van Niel G, D’Angelo G, Raposo G. Shedding light on the cell biology of extracellular vesicles. Nat Rev Mol Cell Biol. 2018;19(4):213–28.

17. Baietti MF, Zhang Z, Mortier E, Melchior A, Degeest G, Geeraerts A, et al. Syndecan–syntenin–ALIX regulates the biogenesis of exosomes. Nat Cell Biol. 2012;14(7):677–85.

18. Ostrowski M, Carmo NB, Krumeich S, Fanget I, Raposo G, Savina A, et al. Rab27a and Rab27b control different steps of the exosome secretion pathway. Nat Cell Biol. 2010;12(1):19–30.

19. Tkach M, Théry C. Communication by Extracellular Vesicles: Where We Are and Where We Need to Go. Cell. 10 mars 2016;164(6):1226–32.

20. Lee YT, Tran BV, Wang JJ, Liang IY, You S, Zhu Y, et al. The Role of Extracellular Vesicles in Disease Progression and Detection of Hepatocellular Carcinoma. Cancers. 2021;13(12):3076.

21. Gest C, Sena S, Dif L, Neaud V, Loesch R, Dugot-Senant N, et al. Antagonism between wild-type and mutant β-catenin controls hepatoblastoma differentiation via fascin-1. JHEP Rep Innov Hepatol. 2023;5(5):100691.

22. Verweij FJ, Bebelman MP, Jimenez CR, Garcia-Vallejo JJ, Janssen H, Neefjes J, et al. Quantifying exosome secretion from single cells reveals a modulatory role for GPCR signaling. J Cell Biol. 2018;217(3):1129–42.

23. Jeppesen DK, Fenix AM, Franklin JL, Higginbotham JN, Zhang Q, Zimmerman LJ, et al. Reassessment of Exosome Composition. Cell. 2019;177(2):428–445.e18.

24. Kowal J, Arras G, Colombo M, Jouve M, Morath JP, Primdal-Bengtson B, et al. Proteomic comparison defines novel markers to characterize heterogeneous populations of extracellular vesicle subtypes. Proc Natl Acad Sci U S A. 2016;113(8):E968–977.

25. Yang L, Peng X, Li Y, Zhang X, Ma Y, Wu C, et al. Long non-coding RNA HOTAIR promotes exosome secretion by regulating RAB35 and SNAP23 in hepatocellular carcinoma. Mol Cancer. 2019;18:78.

26. Boyault S, Rickman DS, de Reyniès A, Balabaud C, Rebouissou S, Jeannot E, et al. Transcriptome classification of HCC is related to gene alterations and to new therapeutic targets. Hepatology. 2007;45(1):42–52.

27. Benhamouche S, Decaens T, Godard C, Chambrey R, Rickman DS, Moinard C, et al. Apc Tumor Suppressor Gene Is the “Zonation-Keeper” of Mouse Liver. Dev Cell. 2006;10(6):759–70.

28. Cadoret A, Ovejero C, Terris B, Souil E, Lévy L, Lamers WH, et al. New targets of β-catenin signaling in the liver are involved in the glutamine metabolism. Oncogene. 2002;21(54):8293–301.

29. Hoshida Y, Nijman SMB, Kobayashi M, Chan JA, Brunet JP, Chiang DY, et al. Integrative Transcriptome Analysis Reveals Common Molecular Subclasses of Human Hepatocellular Carcinoma. Cancer Res. 2009;69(18):7385–92.

30. Shimada S, Mogushi K, Akiyama Y, Furuyama T, Watanabe S, Ogura T, et al. Comprehensive molecular and immunological characterization of hepatocellular carcinoma. EBioMedicine. 2019;40:457–70.

31. Calderaro J, Couchy G, Imbeaud S, Amaddeo G, Letouzé E, Blanc JF, et al. Histological subtypes of hepatocellular carcinoma are related to gene mutations and molecular tumour classification. J Hepatol. 2017;67(4):727–38.

32. Théry C, Witwer KW, Aikawa E, Alcaraz MJ, Anderson JD, Andriantsitohaina R, et al. Minimal information for studies of extracellular vesicles 2018 (MISEV2018): a position statement of the International Society for Extracellular Vesicles and update of the MISEV2014 guidelines. J Extracell Vesicles. 2018;7(1):1535750.

33. Kalra H, Gangoda L, Fonseka P, Chitti SV, Liem M, Keerthikumar S, et al. Extracellular vesicles containing oncogenic mutant β-catenin activate Wnt signalling pathway in the recipient cells. J Extracell Vesicles. 2019;8(1):1690217.

34. Chairoungdua A, Smith DL, Pochard P, Hull M, Caplan MJ. Exosome release of β-catenin: a novel mechanism that antagonizes Wnt signaling. J Cell Biol. 2010;190(6):1079–91.

35. Lu A, Wawro P, Morgens DW, Portela F, Bassik MC, Pfeffer SR. Genome-wide interrogation of extracellular vesicle biology using barcoded miRNAs. eLife. 2018;7:e41460.

36. Lustig B, Jerchow B, Sachs M, Weiler S, Pietsch T, Karsten U, et al. Negative Feedback Loop of Wnt Signaling through Upregulation of Conductin/Axin2 in Colorectal and Liver Tumors. Mol Cell Biol. 2002;22(4):1184–93.

37. Nejak-Bowen K, Monga SP. Wnt/β-catenin signaling in hepatic organogenesis. Organogenesis. 2008;4(2):92–9.

38. Gougelet A, Torre C, Veber P, Sartor C, Bachelot L, Denechaud PD, et al. T-cell factor 4 and β-catenin chromatin occupancies pattern zonal liver metabolism in mice. Hepatol Baltim Md. 2014;59(6):2344–57.

39. Li Z, Fang R, Fang J, He S, Liu T. Functional implications of Rab27 GTPases in Cancer. Cell Commun Signal. 2018;16(1):44.

40. Bobrie A, Krumeich S, Reyal F, Recchi C, Moita LF, Seabra MC, et al. Rab27a supports exosome-dependent and -independent mechanisms that modify the tumor microenvironment and can promote tumor progression. Cancer Res. 2012;72(19):4920–30.

41. Salimu J, Webber J, Gurney M, Al-Taei S, Clayton A, Tabi Z. Dominant immunosuppression of dendritic cell function by prostate-cancer-derived exosomes. J Extracell Vesicles. 2017;6(1):1368823.

42. Marar C, Starich B, Wirtz D. Extracellular vesicles in immunomodulation and tumor progression. Nat Immunol. 2021;22(5):560–70.

43. Berchem G, Noman MZ, Bosseler M, Paggetti J, Baconnais S, Le Cam E, et al. Hypoxic tumor-derived microvesicles negatively regulate NK cell function by a mechanism involving TGF-β and miR23a transfer. Oncoimmunology. 2016;5(4):e1062968.

44. Viel S, Marçais A, Guimaraes FSF, Loftus R, Rabilloud J, Grau M, et al. TGF-β inhibits the activation and functions of NK cells by repressing the mTOR pathway. Sci Signal. 2016;9(415):ra19.

45. Hsu YL, Hung JY, Chang WA, Jian SF, Lin YS, Pan YC, et al. Hypoxic Lung-Cancer-Derived Extracellular Vesicle MicroRNA-103a Increases the Oncogenic Effects of Macrophages by Targeting PTEN. Mol Ther J Am Soc Gene Ther. 2018;26(2):568–81.

46. Park JE, Dutta B, Tse SW, Gupta N, Tan CF, Low JK, et al. Hypoxia-induced tumor exosomes promote M2-like macrophage polarization of infiltrating myeloid cells and microRNA-mediated metabolic shift. Oncogene. 2019;38(26):5158–73.

47. Jullié D, Choquet D, Perrais D. Recycling endosomes undergo rapid closure of a fusion pore on exocytosis in neuronal dendrites. J Neurosci Off J Soc Neurosci. 2014;34(33):11106–18.

48. Sposini S, Jean-Alphonse FG, Ayoub MA, Oqua A, West C, Lavery S, et al. Integration of GPCR Signaling and Sorting from Very Early Endosomes via Opposing APPL1 Mechanisms. Cell Rep. 2017;21(10):2855–67.

49. Bakr M, Jullié D, Krapivkina J, Paget-Blanc V, Bouit L, Petersen JD, et al. The vSNAREs VAMP2 and VAMP4 control recycling and intracellular sorting of post- synaptic receptors in neuronal dendrites. Cell Rep. 2021;36(10):109678.

