## supplemental Figures for "Emerging role of oncogenic β-catenin in exosome biogenesis as a driver of immune escape in hepatocellular carcinoma"

### Figure 1 sup

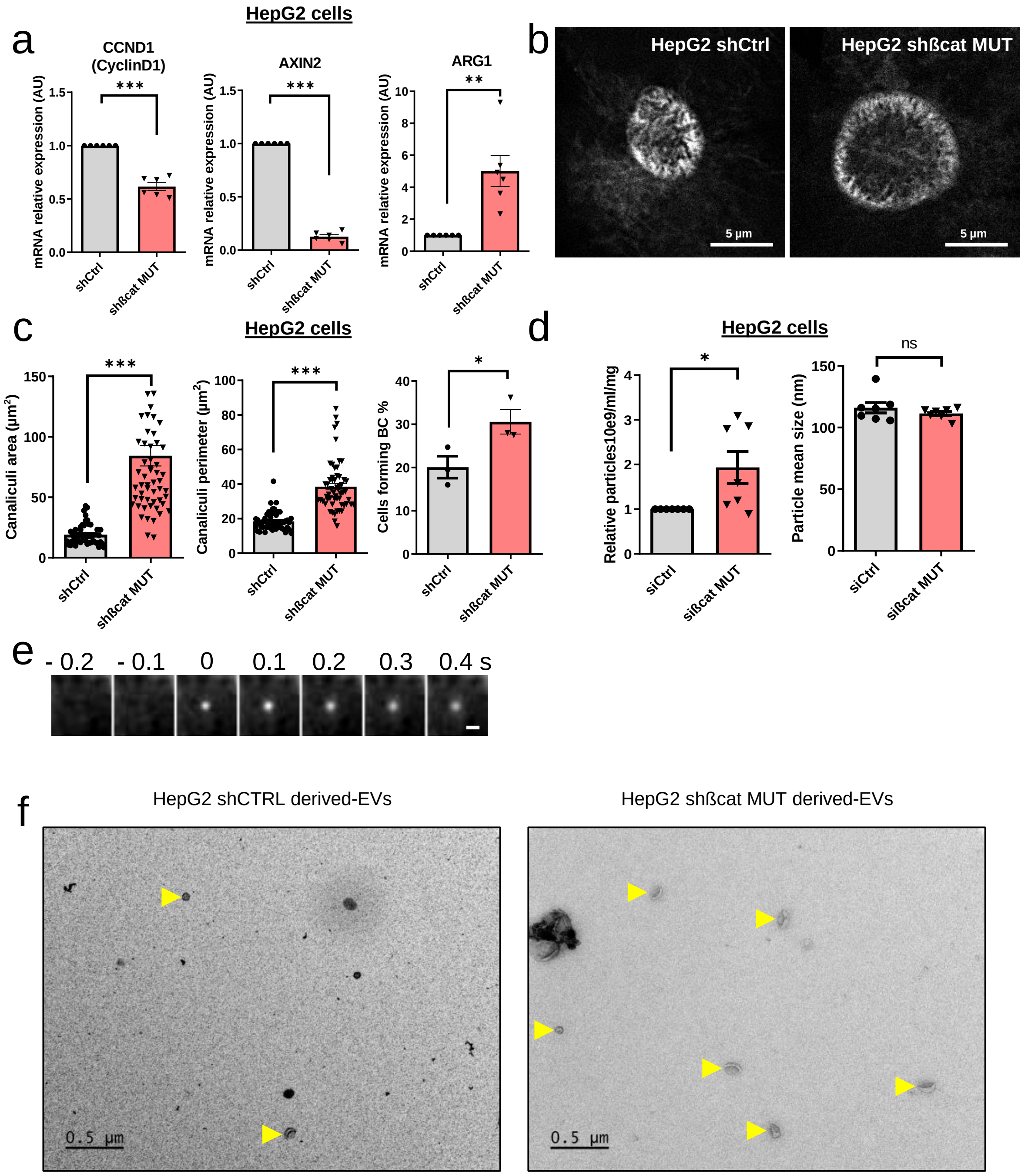
HepG2 cells
a
CCND1 (CyclinD1)
***
mRNA relative expression (AU)
1.5
1.0
0.5
0.0
shCtrl
shßcat MUT
AXIN2
***
ARG1
**
10
8
6
4
2
0
b
HepG2 shCtrl
HepG2 shßcat MUT
5 µm
c
HepG2 cells
Canaliculi area (µm²)
150
100
50
Canaliculi perimeter (µm²)
80
60
40
20
Cells forming BC %
30
*
d
HepG2 cells
Relative particles10e9/ml/mg
Particle mean size (nm)
ns
siCtrl
sißcat MUT
e
- 0.2
- 0.1
0
0.1
0.2
0.3
0.4 s
f
HepG2 shCTRL derived-EVs
HepG2 shßcat MUT derived-EVs
0.5 µm

### Figure 2 sup

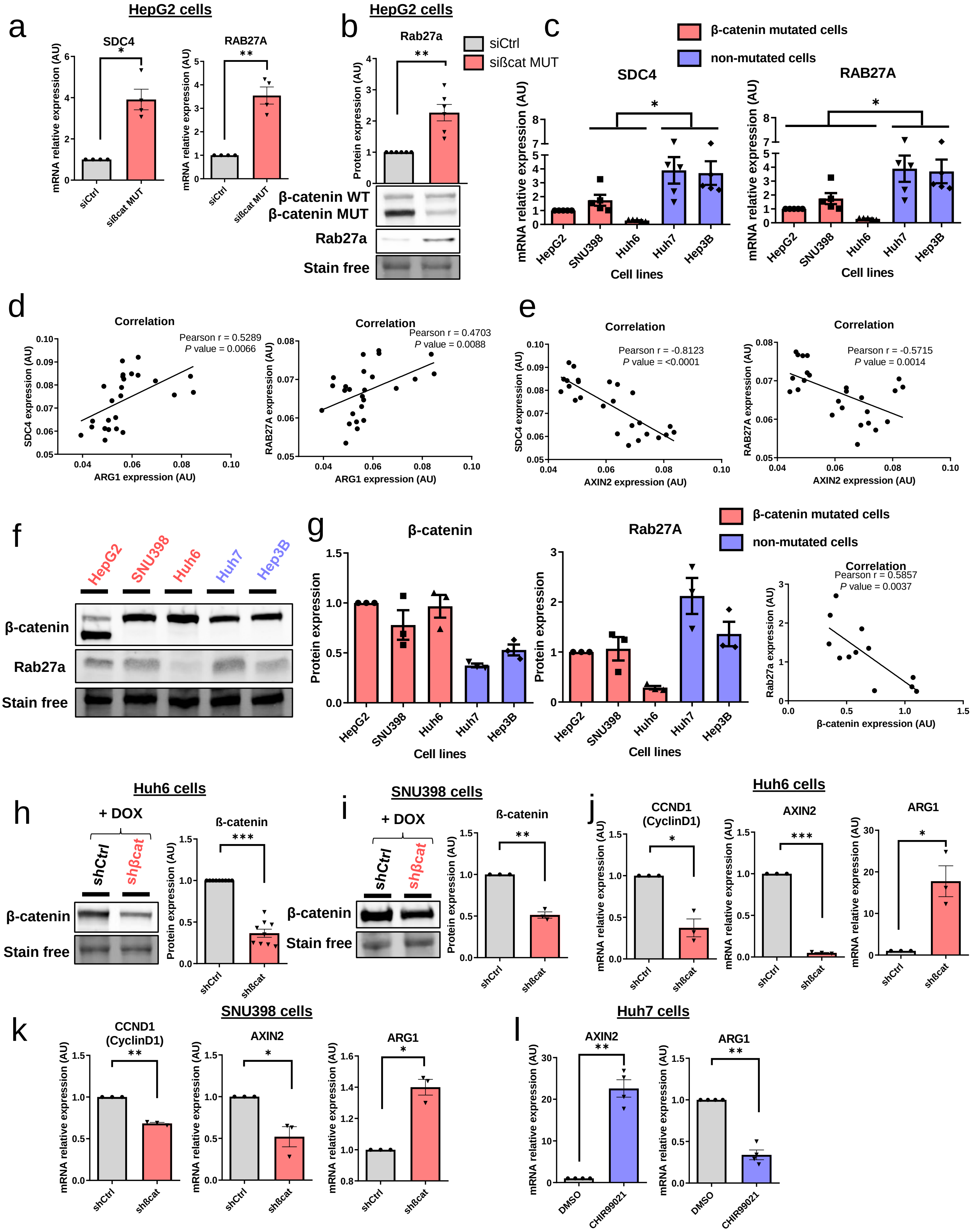

a
HepG2 cells
SDC4
RAB27A
mRNA relative expression (AU)
siCtrl
sißcat MUT
b
HepG2 cells
Rab27a
Protein expression (AU)
siCtrl
sißcat MUT
β-catenin WT
β-catenin MUT
Rab27a
Stain free
c
β-catenin mutated cells
non-mutated cells
SDC4
RAB27A
mRNA relative expression (AU)
HepG2
SNU398
Huh6
Huh7
Hep3B
Cell lines
d
Correlation
Pearson r = 0.5289
P value = 0.0066
SDC4 expression (AU)
ARG1 expression (AU)
Correlation
Pearson r = 0.4703
P value = 0.0088
RAB27A expression (AU)
ARG1 expression (AU)
e
Correlation
Pearson r = -0.8123
P value = <0.0001
SDC4 expression (AU)
AXIN2 expression (AU)
Correlation
Pearson r = -0.5715
P value = 0.0014
RAB27A expression (AU)
AXIN2 expression (AU)
f
HepG2
SNU398
Huh6
Huh7
Hep3B
β-catenin
Rab27a
Stain free
g
β-catenin
Rab27A
Protein expression
Cell lines
β-catenin mutated cells
non-mutated cells
Correlation
Pearson r = 0.5857
P value = 0.0037
Rab27a expression (AU)
β-catenin expression (AU)
h
Huh6 cells
+ DOX
shCtrl
shβcat
β-catenin
Stain free
ß-catenin
Protein expression (AU)
i
SNU398 cells
+ DOX
shCtrl
shβcat
β-catenin
Stain free
ß-catenin
Protein expression (AU)
j
Huh6 cells
CCND1 (CyclinD1)
AXIN2
ARG1
mRNA relative expression (AU)
shCtrl
shßcat
k
SNU398 cells
CCND1 (CyclinD1)
AXIN2
ARG1
mRNA relative expression (AU)
shCtrl
shßcat
l
Huh7 cells
AXIN2
ARG1
mRNA relative expression (AU)
DMSO
CHIR99021

### Figure 3 sup

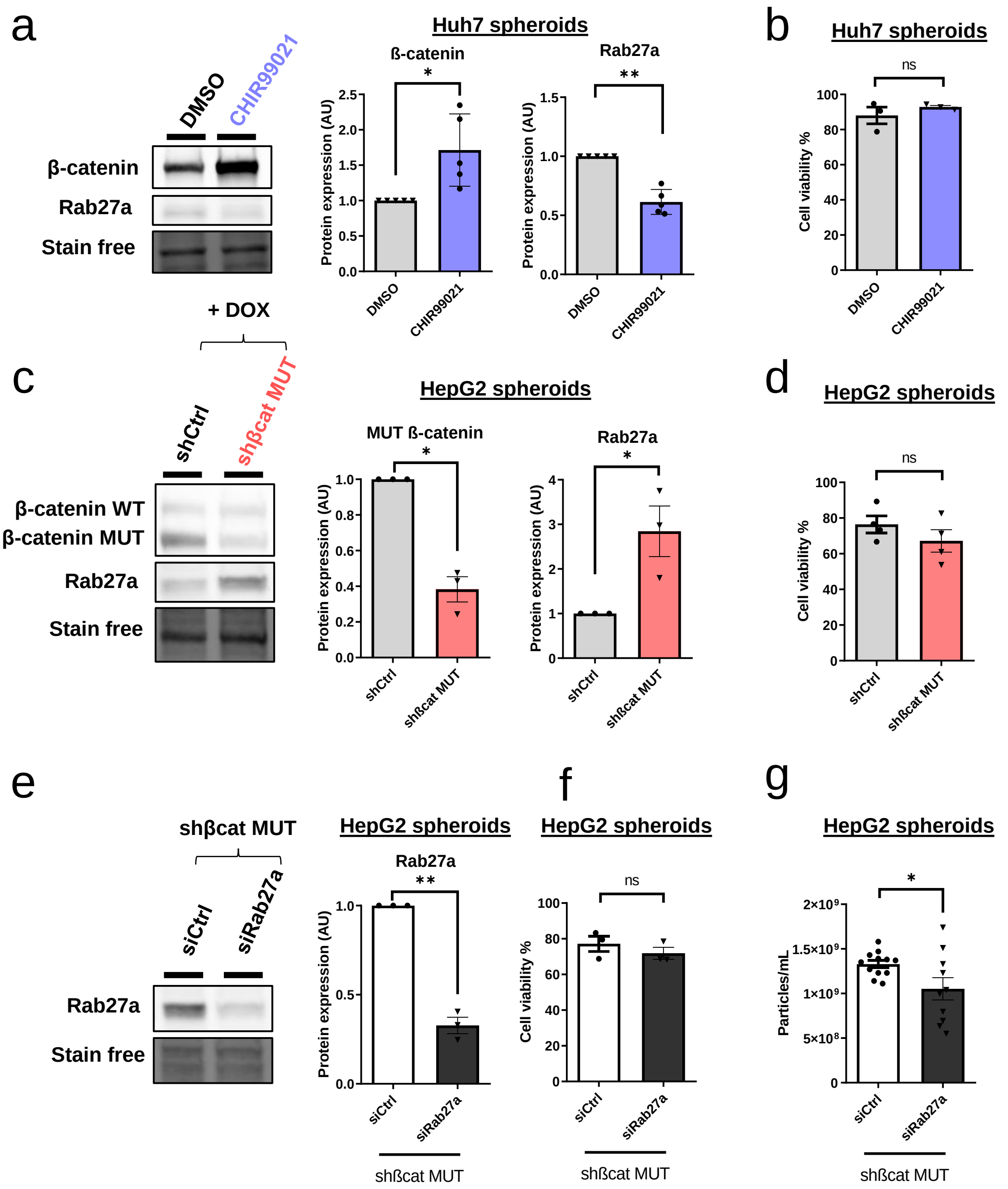
a
Huh7 spheroids
DMSO
CHIR99021
β-catenin
Rab27a
Stain free
ß-catenin
*
Protein expression (AU)
Rab27a
**
b
Huh7 spheroids
ns
Cell viability %
+ DOX
c
HepG2 spheroids
shCtrl
shβcat MUT
β-catenin WT
β-catenin MUT
Rab27a
Stain free
MUT ß-catenin
*
Rab27a
*
d
HepG2 spheroids
ns
Cell viability %
e
shβcat MUT
siCtrl
siRab27a
Rab27a
Stain free
HepG2 spheroids
Rab27a
**
shßcat MUT
f
HepG2 spheroids
ns
Cell viability %
shßcat MUT
g
HepG2 spheroids
*
Particles/mL
shßcat MUT

### Figure 4 sup

a

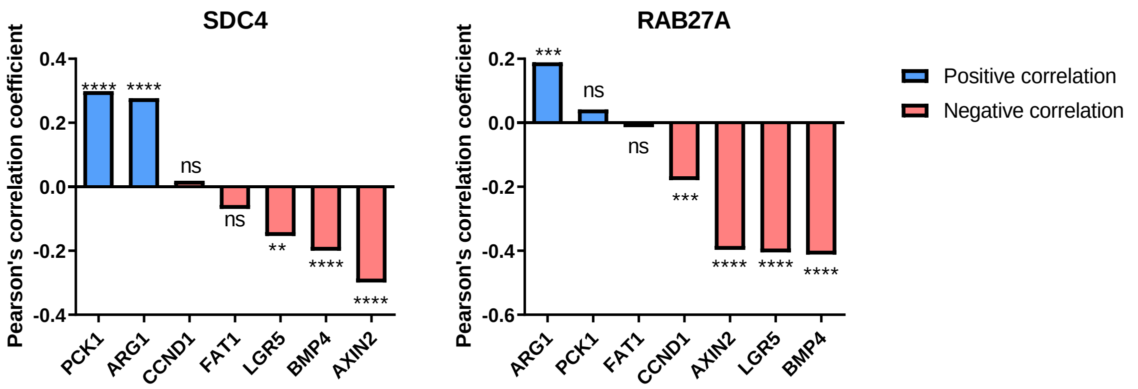


b

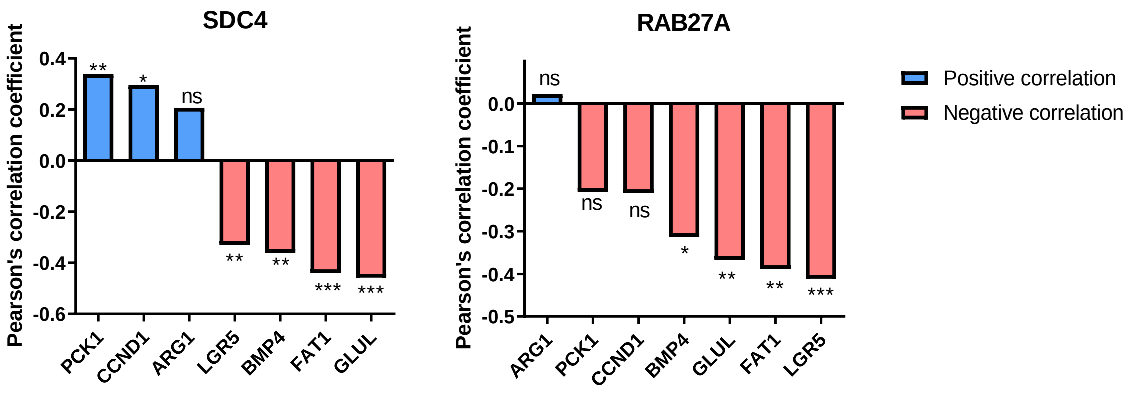


c

| PATIENT COHORT N°1 | |
|---|---|
| **Clinical characteristics** | Value |
| Age (mean ± standard deviation) | 67,9 ± 8,8 years |
| Gender (male) | 94,1% |
| HBV infection | 7,7% |
| HCV infection | 46% |
| Cirrhosis | 50% |
| Metabolic syndrome | 12,5% |
| **Tumor characteristics** | **Value** |
| Diameter (mean ± standard deviation) | 49,9 ± 33,6 |
| Intact capsule | 23,1% |
| Satellite nodule | 50% |
| Vascular invasion | 66,7% |

| PATIENT COHORT N°2 | |
|---|---|
| **Clinical characteristics** | Value |
| Age (mean ± standard deviation) | 69,2 ± 9,7 years |
| Gender (male) | 92,1% |
| HBV infection | 5,4% |
| HCV infection | 14% |
| Cirrhosis | 27% |
| Metabolic syndrome | 14,3% |
| **Tumor characteristics** | **Value** |
| Diameter (mean ± standard deviation) | 68,6 ± 50,5 |
| Intact capsule | 5,4% |
| Satellite nodule | 47,4% |
| Vascular invasion | 70,3% |
